# The glycan shield of alphaherpesvirus glycoprotein B modulates host co-receptor binding

**DOI:** 10.64898/2026.02.03.703472

**Authors:** Sabrina T. Krepel, Francine Rodrigues Ianiski, Bert J.C. Janssen, Joost Snijder

## Abstract

Alphaherpesviruses can infect a wide range of cell types and are notorious for their neurotropism. They enter cells through the fusion activity of the essential viral glycoprotein gB. Interactions of gB with co-receptors Myelin Associated Glycoprotein (MAG), paired immunoglobulin-like type 2 receptor alpha (PILRα) and Non-muscle Myosin Heavy Chain IIA (NMHC-IIA) influence cell tropism and infectivity. Both MAG and PILRα are known to bind to sialic acids, highlighting the potential role for gB glycosylation in mediating the co-receptor interactions. Here, using glycoproteomics, cryo electron microscopy and surface plasmon resonance, we study the specificity of the three co-receptor interactions with gB from herpes simplex virus type 1 (HSV-1), type 2 (HSV-2), and varicella zoster virus (VZV). We show that all gB variants are heterogeneously N-glycosylated at six or seven predicted sites with moderate degrees of sialylation, complemented with extensive O-linked glycosylation at the disordered N-terminus and a flexible loop within domain II. All three gB variants bind to MAG with similar affinity, mediated by sialic acids on gB O-glycans. In contrast, PILRα binds only to HSV-1 and HSV-2, but not VZV gB. PILRα binding is also mediated by sialic acids on gB O-glycans, of which we identify multiple candidate sites in HSV-1 and HSV-2, and none in VZV gB. NMHC-IIA binds to HSV-1 gB and HSV-2 gB with high affinity through a glycan-independent interaction, and not to VZV gB. These data reveal divergent co-receptor specificity and interaction strength and provide an improved biophysical basis to understand human neurotropic herpesvirus infection.

## Introduction

The human alphaherpesviruses Herpes Simplex Virus type 1 (HSV-1), type 2 (HSV-2) and Varicella Zoster Virus (VZV) can infect a wide range of human tissues and are notorious for their neurotropism. The effect of neuron invasion can lead to directly associated diseases such as Herpes Simplex Encephalitits (HSE)^1,2^. More recently, correlations between increased herpesvirus titers in the brain and various well-known neurological diseases like dementia, multiple sclerosis, and Alzheimer’s disease have been reported as well^3–6^. Glycoprotein B (gB) on the surface of herpesviruses plays an important role in the infection of mammalian cells as a fusogen. While the main host receptor interactions are mediated by herpesvirus glycoproteins gH/gL and gD, gB is also known to bind co-receptors to increase infectivity and thereby shape tropism of the virus^7,8^. Three co-receptors have been identified for glycoprotein B: Myelin-Associated Glycoprotein (MAG), paired immunoglobulin-like type 2 receptor alpha (PILRα) and Non-muscle Myosin Heavy Chain IIA/B (NMHC-IIA/B) that enhance cell infection.

The roles of MAG, PILRα and NMHC-IIA in herpesvirus infection vary. For HSV-1 and VZV, infectivity increases through gB-MAG association in MAG-transfected cells^9^. For VZV, MAG-mediated infection is induced by gB and gH/gL only, whereas for HSV-1, gD is an additional requirement, indicating that MAG-mediated infection remains dependent on initial interaction of gD with host receptor Nectin-1^7,10^. Interestingly, PILRα expression on CHO-cells deficient of Nectin-1 enables HSV-1 infection but not HSV-2 infection, although a slight increase of HSV-2 infection is observed when Nectin-1 is present^11–14^. The development of herpes keratitis following corneal infection is dependent on the ability of HSV-1 gB interacting with PILRα. Moreover, brain tissue with low gD receptor expression have high PILRα expression and are also vulnerable to infection^15^, suggesting that the gB– PILRα interaction plays a role in the brain. The infection rate of HSV-1 increases when NMHCII-A/B are overexpressed in Vero or IC21 cells and decreases when expression is knocked down^16–18^. While NMHC-IIA/B typically localize intracellularly, trafficking of NMHC-IIA/B to the cell surface has been reported^16^, and cells infected with HSV-1 increase cell surface presentation of NMHCII-A. In addition, interaction of the transmembrane protein TMEFF1 with Nectin-1 and NMHC-IIA/B may protect against HSV-1 infection in the central nervous system (CNS).^19^ Besides the extracellular role as a host receptor, NMHC-IIA has also been suggested as an intracellular receptor. Walking of the myosin NM-IIA – consisting of NMHC-IIA – towards the minus end of actin filaments creates pulling forces that lead to virus entry^20^.

The three gB receptors have differences in structure, function, and expression. MAG, also known as Siglec4, is mainly expressed in the nervous system on the surface of oligodendrocytes forming myelin-axon sheets, where it maintains the correct sheet distance to the axon by interacting with α2,3-linked sialosides of neuronal gangliosides^21,22^. MAG consists of 5 Ig-like domains, of which the interface between domain 4 and 5 establishes a dimeric state and domain 1 interacts with its ligand^23^. It is regarded as a key protein in inhibition of axonal regeneration after neuronal injury in the CNS^24^. A structurally similar protein, PILRα, is mainly expressed by immune cells, including microglia in the brain^25^. PILRα acts as an opposite equivalent of PILRβ by inhibiting immune cell activation through integrin modulation and modulates neutrophil infiltration and cytokine release during inflammation ^26,27^. Significantly smaller than MAG, the extracellular domain of PILRα adopts a V-set Ig-like fold which can bind to various ligands such as CD22 and NPDC1, mediated by sialylated O-glycans and nearby ligand residues^28–31^. NMHC-IIA and NMHC-IIB, also known by their gene names *myh9* and *myh10*, are isoforms of a subunit belonging to non-muscle myosin IIA (NM-IIA) and are structurally unrelated to the receptors MAG and PILRα. NMHC-IIA is involved in cellular integrity, cell adhesion and filopodia formation and is ubiquitously expressed in human tissues with heavy chain isoform expression divergence^32–34^. Although isoform NMHC-IIA is primarily expressed in epithelial cells, NMHC-IIB is mainly found in neurons. These molecules consist of two major domains: a motor domain followed by a 1123-amino acid alpha helix that forms a coiled-coil upon dimerization. The differences in structure suggest that the three host receptors differ in their mode of interaction with the gB proteins of Herpesviruses.

The molecular interplay of gB receptors and their contribution to neuroinvasion is poorly understood. Differences between host co-receptor and HSV-1, HSV-2 and VZV gB interactions have been reported before. Sialosides on VZV gB are crucial for MAG-gB interaction, which is further supported by the inability of VZV gB to bind to MAG^R118A^, a mutant that is unable to bind to sialic acids^35^. The VZV gB-MAG interaction is not diminished in alanine scanning mutagenesis of a wide range of singular putative O-linked and N-linked sites in gB, indicating that the interaction is promiscuous and not pinned down to a single specific glycosylation site ^35^. In contrast, functional interaction of PILRα with HSV-1 gB was reported to depend on two specific O-glycan sites, predominantly T53 and to a lesser degree T480^36^. Crystal structures and binding assays with synthetic O-glycopeptides have revealed that PILRα binds to α2,6-linked sialic acids on the core O-linked N-Acetylgalactosamine and makes additional contacts with the carbonyl backbone and pyrrolidine ring of an adjacent proline residue at the +1 position, relative to the T53 site of O-glycosylation in HSV-1 gB. While this proline is also present in other known O-linked glycoprotein interaction sites of PILRα, it remains unclear whether this +1 proline is a strict requirement for binding^29,30,36–38^. Interestingly, HEK293T cells expressing VZV gB do not interact with a dimerized construct of PILRα^39^, which suggests that VZV gB lacks the required O-linked α2,6-sialoside or the putative associated +1 proline motif. Few studies have investigated the molecular mechanism of NMHCII-A/B binding to gB, although it has been reported that the C-terminal coiled coil region of NMHCII-A/B interacts with gB^16^. These previous studies indicate differences in specificity of the three gB proteins and the three receptors, but information on the affinities and binding sites of these interactions is lacking.

In this study, we assess the binding affinities between HSV-1, HSV-2 and VZV gB and their receptors MAG, PILRα and NMHCII-A. We find a hierarchy of interactions and the presence of multiple interactions sites on the gB proteins. We determined binding affinities by quantitative *in vitro* binding assays, map candidate binding sites with glycoproteomics and determined the structural context of these glycan-mediated interactions in a 2.8 Å cryo-EM structure of unliganded HSV-2 gB in its postfusion conformation. Our data show that all three gB variants interact with MAG via their sialosides of predominantly O-glycans, whereas only HSV-1 and HSV-2 gB bind to PILRα. Mutation of two highly occupied O-glycan sites with +1 proline residues in HSV-2 gB did not abrogate binding, suggesting the presence of additional PILRα binding sites and promiscuous interaction with O-linked sialylated O-glycans in HSV-2. HSV-1 and HSV-2 gB bind to NMHCII-A with substantially higher affinity than to MAG and PILRα. This interaction is not affected by N-glycan modifications, providing first evidence for protein-protein mediated interactions of gB with a co-receptor. VZV gB did not associate with PILRα nor with NMHCII-A, of which the former is explained by the lack of O-glycan sialosides with a putative PILRα binding motif. These findings reveal unexpected differences and similarities between HSV-1, HSV-2, and VZV gB-receptor interactions.

## Results

### N-linked glycans in alphaherpesvirus are heterogeneous and sparsely sialylated

Previous reports highlight that MAG and PILRα bind to gB via N– and/or O-linked sialosides. To gain more insight into the type and location of N– and O-linked glycans presented on gB variants, and pinpoint candidate interaction sites with MAG and PILRα, we performed a comparative glycoproteomics analysis of gB ectodomains in their postfusion state. We compared gB constructs from HSV-1, HSV-2, and VZV, which we will refer to as gB1, gB2 and gBV, respectively. The sequences of gB1, gB2, and gBV carry 6, 7, and 7 predicted N-linked glycosylation sequons (NXS/T, X≠P), respectively (Fig. 1A). Of those, three sites are fully conserved across all gB variants: N141, N430, and N674 (following gB1 numbering). In addition, site N87, N398 and N489 are also conserved between gB1 and gB2, the latter containing one unique site at N473. The gBV sequence contains 4 unique sites at N257, N479, N503, and N620.

**Figure 1.**
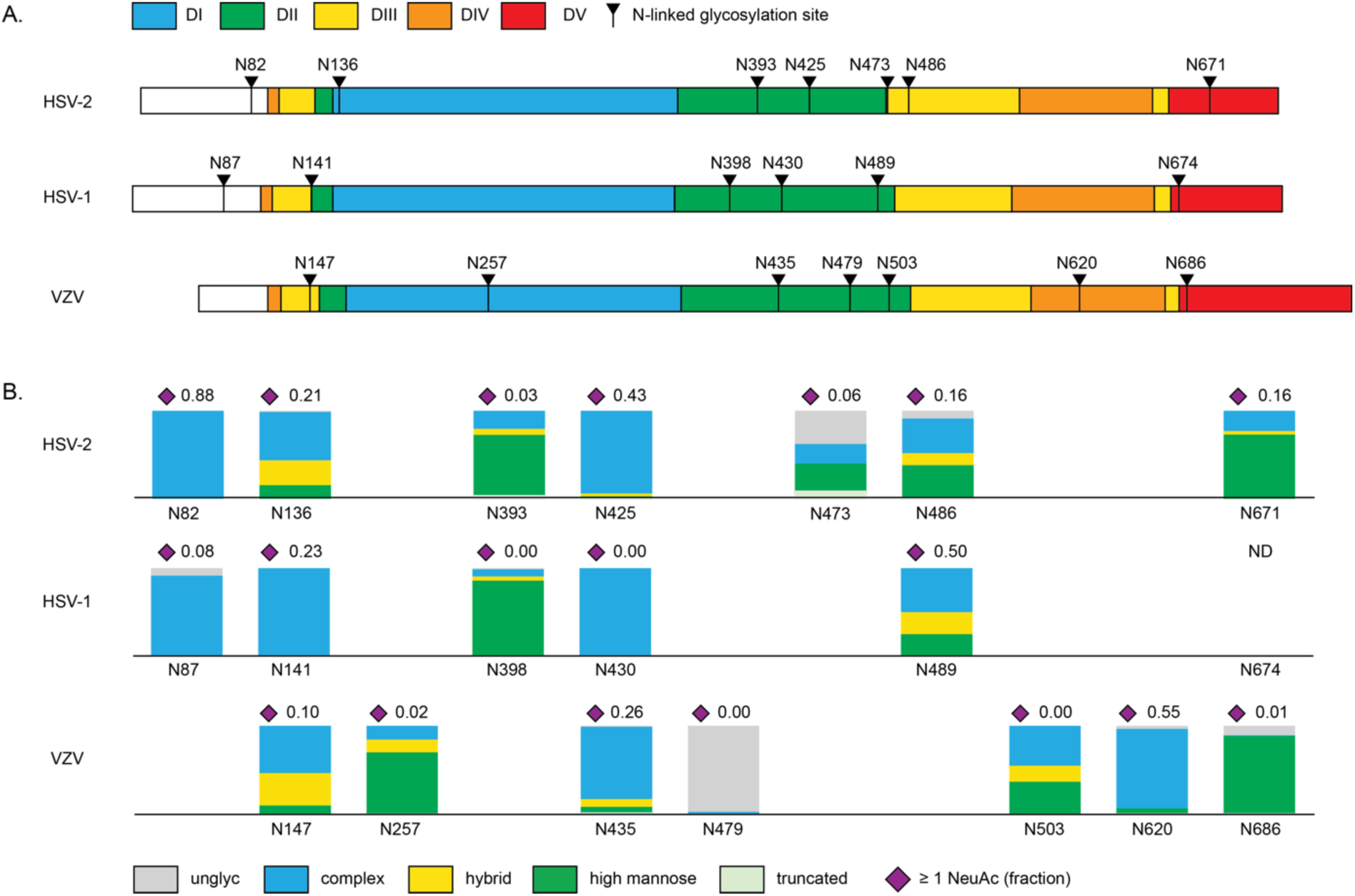
N-linked glycosylation of gB. A. Domain structure of the three gB variants, coloured according to Grunewald et al^40^ with predicted N-linked glycosylation sites. B. N glycoproteomics analysis of gB variants. Plotted is the fraction of identified glycopeptide spectra categorized as high-mannose (=2 HexNAc), hybrid (=3 HexNAc), or complex (≥4 HexNAc), in addition to truncated (core pentasaccharide or smaller), and unglycosylated (unmodified asparagine), coloured following the Symbol Nomenclature For Glycans. The fraction of glycopeptides with at least 1 NeuAc residue is plotted above each site, indicated with the purple diamond.

The identified N-glycopeptides in our proteomics analyses cover all sites in each gB variant, except for N674 in gB1, which remains a blind spot in this experiment (Fig. 1B and Supplementary Data S1). As expected with protein produced in HEK293 cells, all sites in the three gB variants display a heterogenous mixture of glycan compositions, dominated by complex di– and tri-antennary glycans overall, but some selected sites stand out with signs of unprocessed high-mannose glycans or even unmodified free asparagines. Specifically, the conserved N674 site in domain V carries predominantly high-mannose glycans. Likewise, N398 in domain II, which is shared between gB1 and gB2, shows a consistent elevated level of high-mannose glycans in both gB variants. The same is seen for the unique site N257 of gBV. Notably, the unique site N473 of gB2 additionally shows a high degree of unmodified asparagine and the unique site N479 of gBV even appears to remain almost completely unmodified. While complex glycans are present at the majority of N-linked sites, they appear only sparsely sialylated, further emphasizing the potential role for O-linked glycans in mediating the MAG and PILRα interactions.

### O-linked glycans in gB present multiple candidate binding sites for MAG and PILRα

We extended our glycoproteomics analyses to O-linked glycosylation by repeating the LC-MS/MS experiments following PNGase F digestion of gB. Enzymatic release of N-glycans by PNGase F converts N-linked sites to deamidated asparagines (*i.e.* aspartic acids with a +1 Da mass shift relative to unmodified asparagine). Removal of N-linked glycans simplifies glycoproteomics analysis of O-linked sites, while retaining information on the glycosylation status of adjacent N-linked sites. We identified a multitude of O-linked glycosylation sites in all three gB variants, predominantly in the disordered N-terminus and a flexible loop in domain II (Fig. 2 and Supplementary Data S2). The identified O-linked glycans are heterogeneous but dominated by core 1 (T-antigen), sialyl core 1, and disialyl core 1 structures.

**Figure 2.**
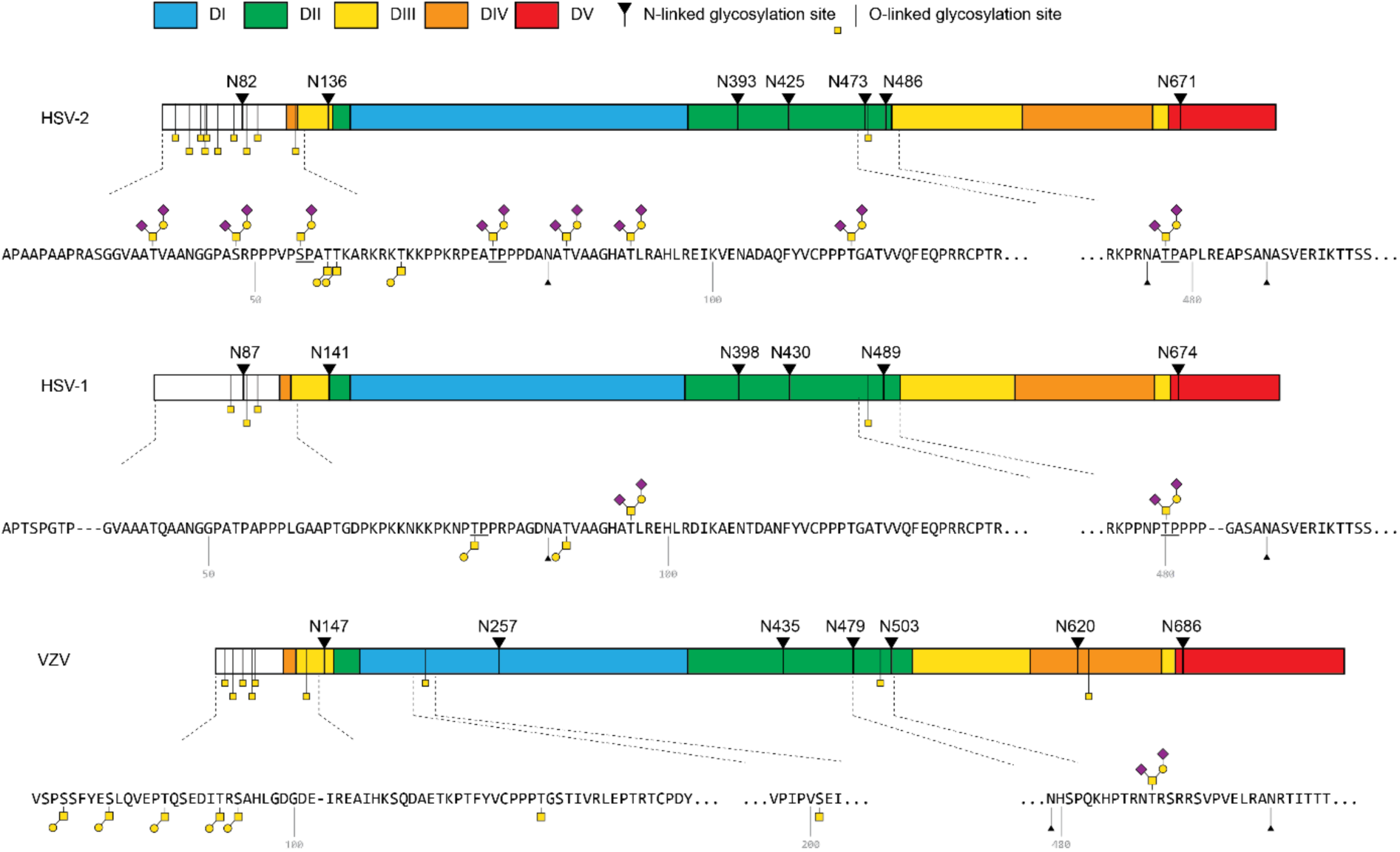
O-linked glycoproteomics of gB variants. Shown are the domain structures of gB variants, with predicted N-glycosylation sites, and experimentally determined O-linked sites. O-linked sites are indicated with yellow squares. The highlighted sequences plot the most frequently identified O-glycoform following standard Symbol Nomenclature For Glycans: GalNAc yellow square, Gal yellow circle, NeuAc purple diamond. Potential PILRα binding sites with proline at the +1 position

Interestingly, the α2,3-linked NeuAc in sialyl core 1 structures are potential ligands for MAG binding, and with the additional α2,6-linked NeuAc in disialyl core 1, such O-linked glycans may double as ligand for both MAG and PILRα. Following this binding specificity, we identify two candidate binding sites for MAG and PILRα in gB1, of which only T480 has the +1 proline of the putative PILRα binding motif. In gB2, we identified 9 candidate binding sites, of which S55, T76, and T475 carry the +1 proline motif. For gBV, we identified only two sialylated O-linked glycosylation sites (T85 and T486), of which T85 carried predominantly unsialylated core 1 structures, and neither carry the +1 proline motif. Of note, the previously identified PILRα binding site T53 in gB1 was not observed to contain sialylated O-linked glycans in our experiments. In gB2, the candidate PILRα binding site T475 is conserved and functionally relevant in gB1 (T480), but the S55 and T76 sites are novel. Based on glycopeptide spectral counts, the O-linked glycosylation of T76 and T475 in gB2 appears to be especially extensive. Interestingly, T475 in gB2 is part of an N-linked glycosylation sequon. Detection of deamidated asparagine in the identified O-glycopeptides shows that both modifications co-exist on the same polypeptide chain, suggesting possible crosstalk between the two types of glycosylation. A second O-linked site within an N-linked sequon is also found at T84 in gB2 and the equivalent T89 in gB1. We and others have previously reported on such combined O-linked glycosylation in N-linked sites for coronavirus Spike proteins, ebolavirus glycoprotein, as well as a multitude of secreted and transmembrane human glycoproteins^41–44^.

### Binding sites for MAG and PILRα on gB are clustered at domain II and the disordered N-terminus

To place the observed glycosylation patterns into their 3D structural context, we determined the structure of postfusion gB2 to 2.8 Å resolution by cryogenic electron microscopy (Fig. 3, Suppl. Fig. S5, Suppl. Tab. S1). While working on the current study other postfusion gB2 structures were published^45–47^, yielding virtually identical atomic models of the postfusion trimer. The ordered structure of postfusion gB2 spans domains I-V, with the disordered N-terminus (residues 31-98) and a flexible loop in domain II (residues 461-486) not visible in the map. This structurally ordered part of postfusion gB2 contains N-glycans at positions N136, N393, N425 and N671, which all show clear low-resolution density for core N-acetylglucosamine moieties and are accessible on the surface of gB2. The remaining three glycosylation sites, N82 in the disordered N-terminus and N473-N486 in the flexible loop of domain II, are not visible in the map but must clearly be exposed and available for co-receptor binding. We observed no visible density for any of the experimentally determined O-linked glycosylation sites, which, except for T115 in domain IV, are all present in the disordered N-terminus and flexible loop of domain II.

**Figure 3.**
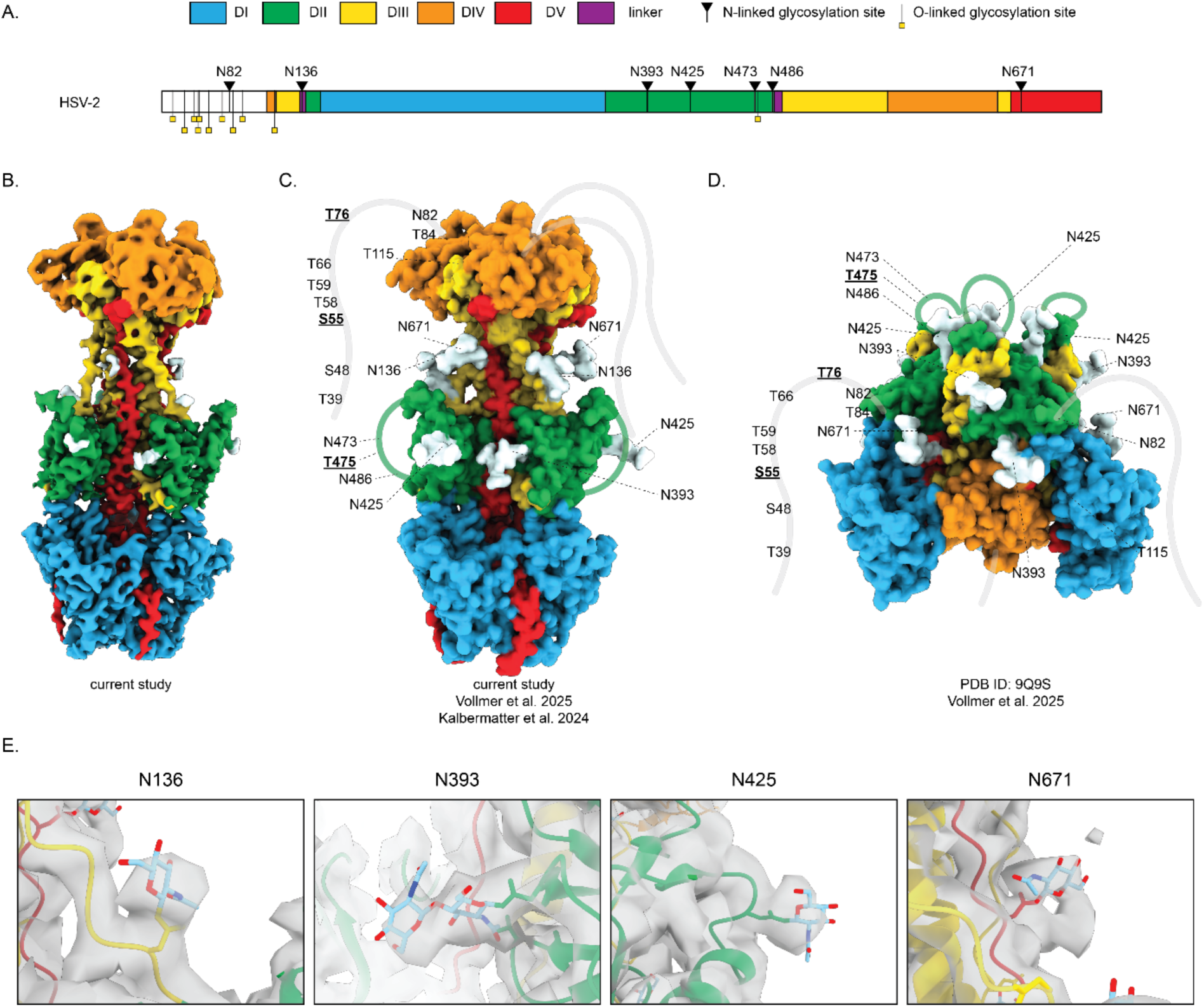
Structural context of gB2 glycosylation. A. Domain structure of gB2 with indicated N– and O-glycosylation sites. B. Experimentally determined single particle cryoEM structure (2.8 Å) density map of postfusion gB2 ectodomain. C. Surface rendering of the postfusion gB2 atomic model built in the cryoEM map, with core pentasaccharide N-glycans grafted on the model displayed in white. Recently determined similar structures are referenced below^45,48^. D. Surface rendering of prefusion gB2 atomic model recently reported by Vollmer et al^45^. E. Glycan cores visualized in the cryo-EM map of gB2. Only glycans showing density are modeled.are underlined.

While the N-linked glycosylation sites in the ordered regions of postfusion gB2 are spread across domains II, III and V, they all fold together at the waist region of the postfusion stalk, concentrated at domain II. Likewise, in the recently reported prefusion structure of gB2^45^, the glycosylation sites within the ordered parts cluster together at domain II, now presented at the apex of the prefusion stump. Taken together, our structural and glycoproteomics analyses suggest that MAG may bind gB at any of the sparsely sialylated N-glycans distributed across the disordered N-terminus and domains II, III and V, in addition to the multitude of O-linked sites. With exception of N87 in the disordered N– terminus, most N-linked sites cluster together at the waist region of the postfusion gB conformation, or the apex of its prefusion state. In contrast, potential binding sites for PILRα are restricted to the disordered N-terminus and the flexible loop of domain II in gB1 and gB2.

### MAG binds indiscriminately to all gB variants

We characterized the glycan-dependent interactions between gB1, gB2, gBV and their host co-receptors MAG, PILRα and NMHC-IIA with SPR (see Fig. 4 and Table 1 for an overview). All three gB’s bind to MAG with similar affinities (KD 25.1 ± 0.7 µM, 35.1 ± 1.2 µM, 24.9 ± 3.5 µM, for gB1, gB2 and gBV, respectively, Suppl. Fig. S1A). Binding of gB1, gB2 and gBV to MAG with low micromolar affinity is typical of MAG’s sialic acid binding. Indeed, when we tested binding of mutant MAG^R118A^, which is deficient in sialic acid binding by removing a key electrostatic interaction^23^, no binding was observed (Fig. 5A-C).

**Figure 4.**
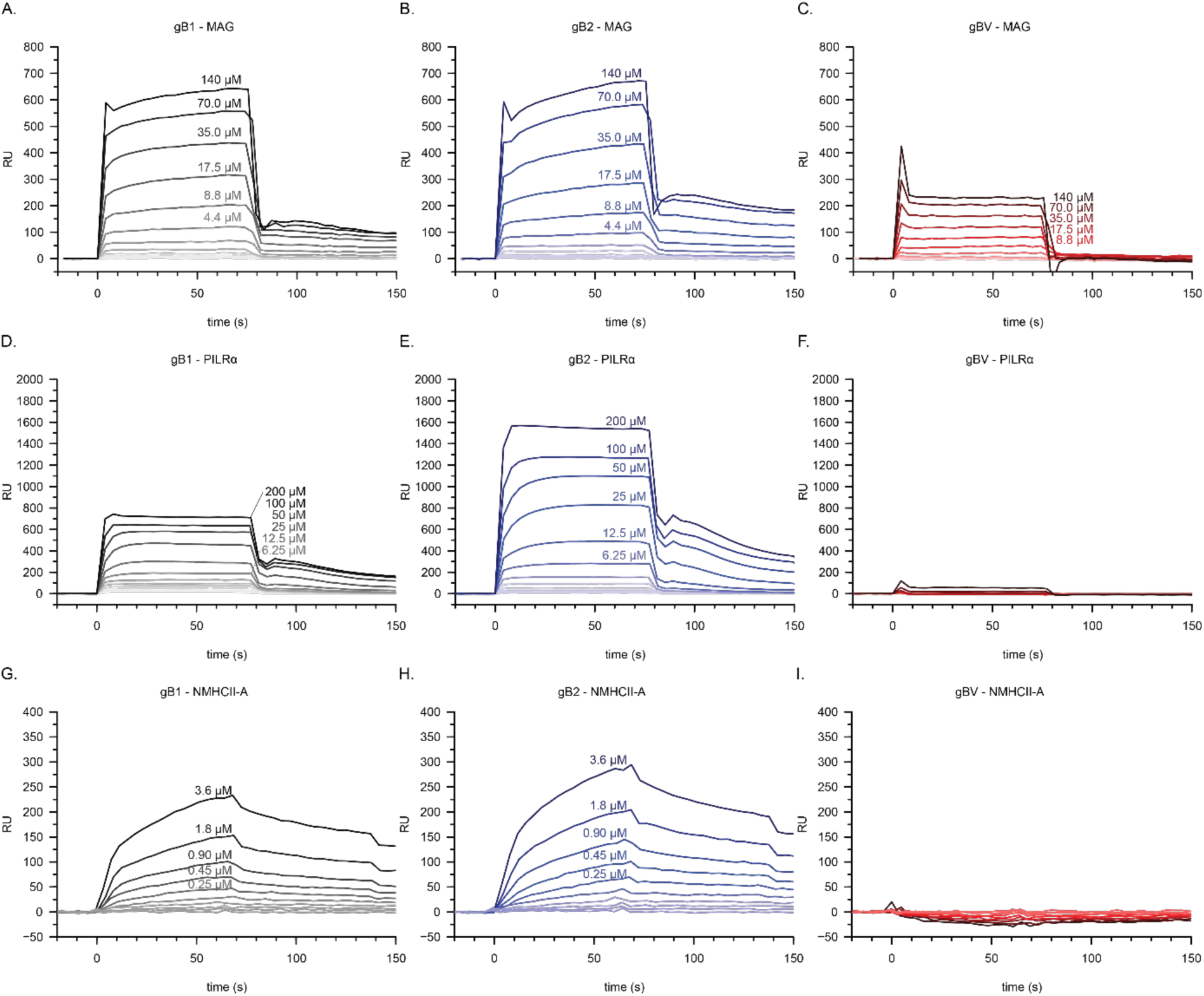
Interactions of three herpesvirus gB variants with three host co-receptors MAG, PILRα and NMHCII-A as determined by SPR. A-I. SPR sensograms with the concentrations of analytes indicated. In each experiment a gB was coupled to the SPR surface. No binding was observed for gBV to PILRα (F) and NMHC-IIA (I).

**Figure 5.**
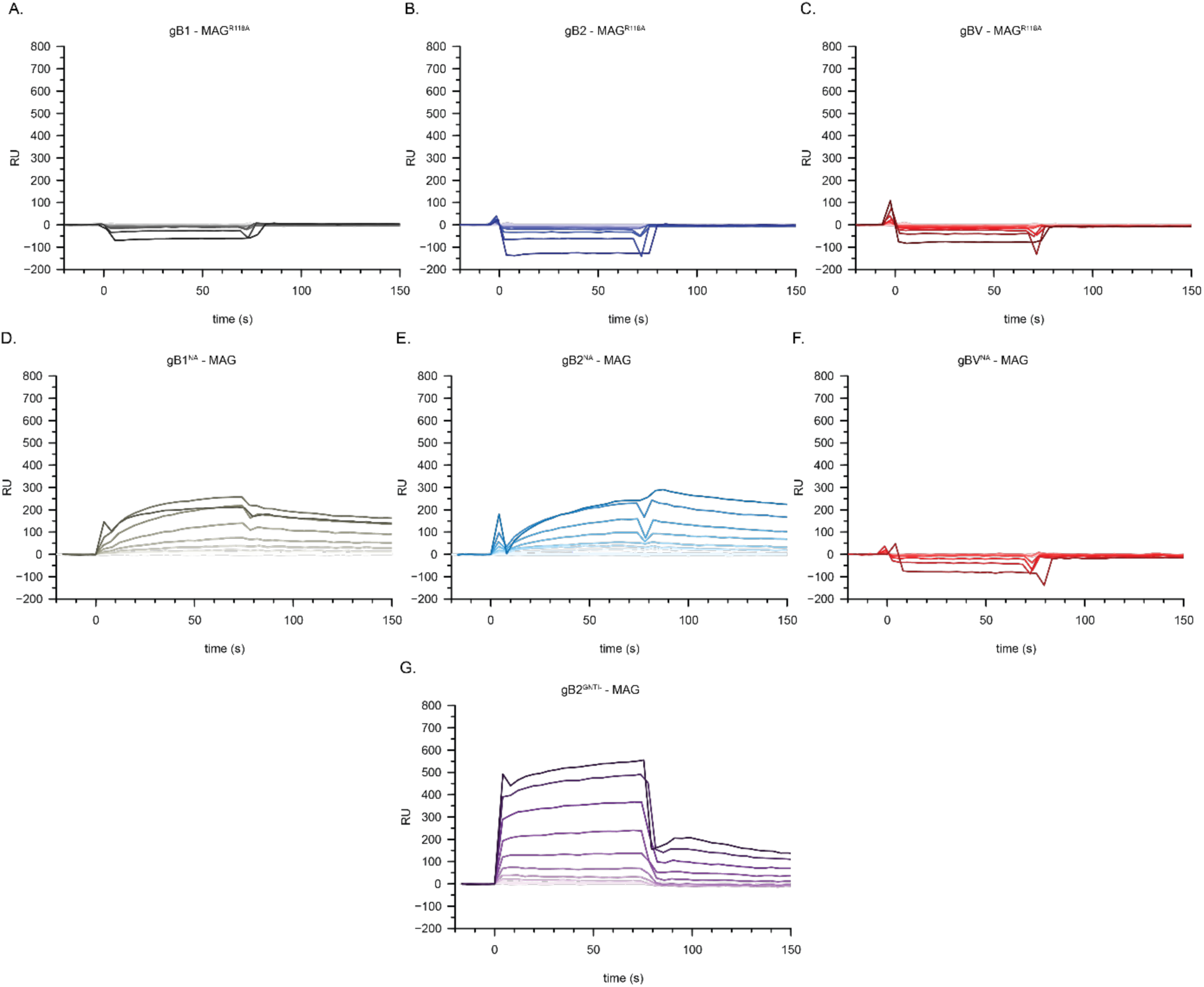
Sialic acid dependence of MAG-gB interaction. SPR sensograms are shown. In each experiment gB was coupled to the SPR surface. A-C. MAG^R118A^ (two-fold dilutions, maximum concentration 140 μM), deficient in sialic acid binding, does not interact with any of the gB’s. D-F. Neuraminidase (NA) treated gB’s have reduced binding to MAG. G. Interaction is unacected of MAG binding to gB2 containing sialylated O-linked glycans but lacking sialylated N-linked glycans. In D-G, MAG at two-fold dilutions, maximum concentration 140 μM.

**Table 1.**
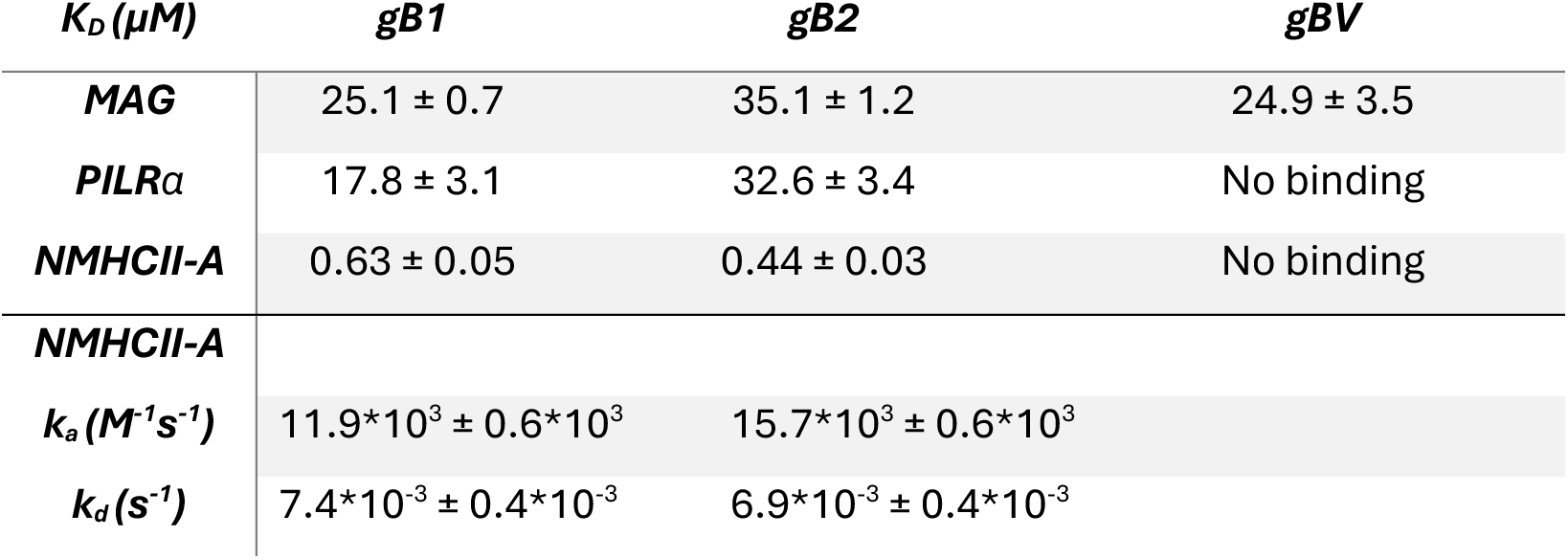
K_D_ values calculated for each gB-receptor interaction. For MAG and PILRα, SPR equilibrium binding plots were fitted to a 1:1 Langmuir binding model (Suppl. Fig. S1) to calculate K_D_. For NMHCII-A, SPR sensograms were fitted to a 1:1 kinetic model to determine k_a_, k_d_ and K_D_. (Suppl. Fig. S3).

| $K_D$ (μM) | gB1 | gB2 | gBV |
| --- | --- | --- | --- |
| **MAG** | 25.1 ± 0.7 | 35.1 ± 1.2 | 24.9 ± 3.5 |
| **PILRα** | 17.8 ± 3.1 | 32.6 ± 3.4 | No binding |
| **NMHCII-A** | 0.63 ± 0.05 | 0.44 ± 0.03 | No binding |
| **NMHCII-A** |  |  |  |
| **$k_a$ ($M^{-1}s^{-1}$)** | 11.9*10<sup>3</sup> ± 0.6*10<sup>3</sup> | 15.7*10<sup>3</sup> ± 0.6*10<sup>3</sup> | |
| **$k_d$ ($s^{-1}$)** | 7.4*10<sup>-3</sup> ± 0.4*10<sup>-3</sup> | 6.9*10<sup>-3</sup> ± 0.4*10<sup>-3</sup> | |

In addition, enzymatic trimming of sialic acids on gB reduced binding, albeit not completely for gB1 and gB2, likely due to incomplete digestion (Fig. 5D-F). Repetition of this experiment for gB2, using fresh neuraminidase, completely abolished binding of MAG (Suppl. Fig. S2), further confirming that the gB-MAG interaction is indeed mediated by sialiosides presented on the gB surface. gB2 produced in HEK293 GnTI−/− cells, a knockout that blocks processing of oligomannose N-linked glycans into complex sialylated structures, binds to MAG with a KD of 34 µM (Fig. 5G), i.e. the same affinity as MAG binding to gB2 containing complex sialylated structures. This indicates that sialylated N-glycans are not essential in MAG-gB binding, and that sialylated O-linked glycans can fully account for the interaction instead.

### PILRα binds to gB1 and gB2, but not gBV

In a similar quantitative interaction experiment, replacing analyte MAG with PILRα, we observe a comparable affinity of PILRα binding to gB1 and gB2 (K_D_ 17.8 ± 3.1 µM and 32.6 ± 3.4 µM, respectively, Fig. 4D-F, Suppl. Fig. S1B). In contrast to MAG, PILRα does not bind to gBV. This is in accordance with our glycoproteomics data that indicates PILRα binding sites are absent in gBV (Fig. 2). gB2 expressed by GnTI−/− cells, has a similar affinity for PILRα compared to gB2 containing complex N-linked glycans (Fig. 6A), supporting the previously reported specificity of PILRα to O-glycans^29^. For HSV-1, it was previously reported that mutation of O-glycan sites T53 and T480 results in diminished association with PILRα. Of these, our glycoproteomics data show that only gB1 T480 contains disialyl core 1 linked glycans (Fig. 2). The equivalent site in gB2, T475, also has disialyl core 1 glycans. Based on the counts of identified glycopeptides, gB2 T475 together with T76 appear to be especially extensively glycosylated. This, together with the conservation of T475, numbered T480 in gB1, led us to test PILRα binding of a gB2 double mutant, T76A/T475A. We find that mutant gB2^T76A/T475A^ is still able to bind to PILRα, albeit with a slightly lower affinity (i.e. a K_D_ of 32.6 µM for wt gB2 versus 42.2 µM for gB2^T76A/T475A^, Fig. 6B). This suggests the existence of additional binding sites for PILRα in gB2, most likely S55 (with the +1 proline motif) identified in our glycoproteomics experiments (Fig. 2). The lack of gBV binding to PILRα indicates that required interaction sites are absent, in concordance with our glycoproteomics results (Fig. 2), and shows that MAG is a more promiscuous receptor for herpesvirus gB’s compared to PILRα.

**Figure 6.**
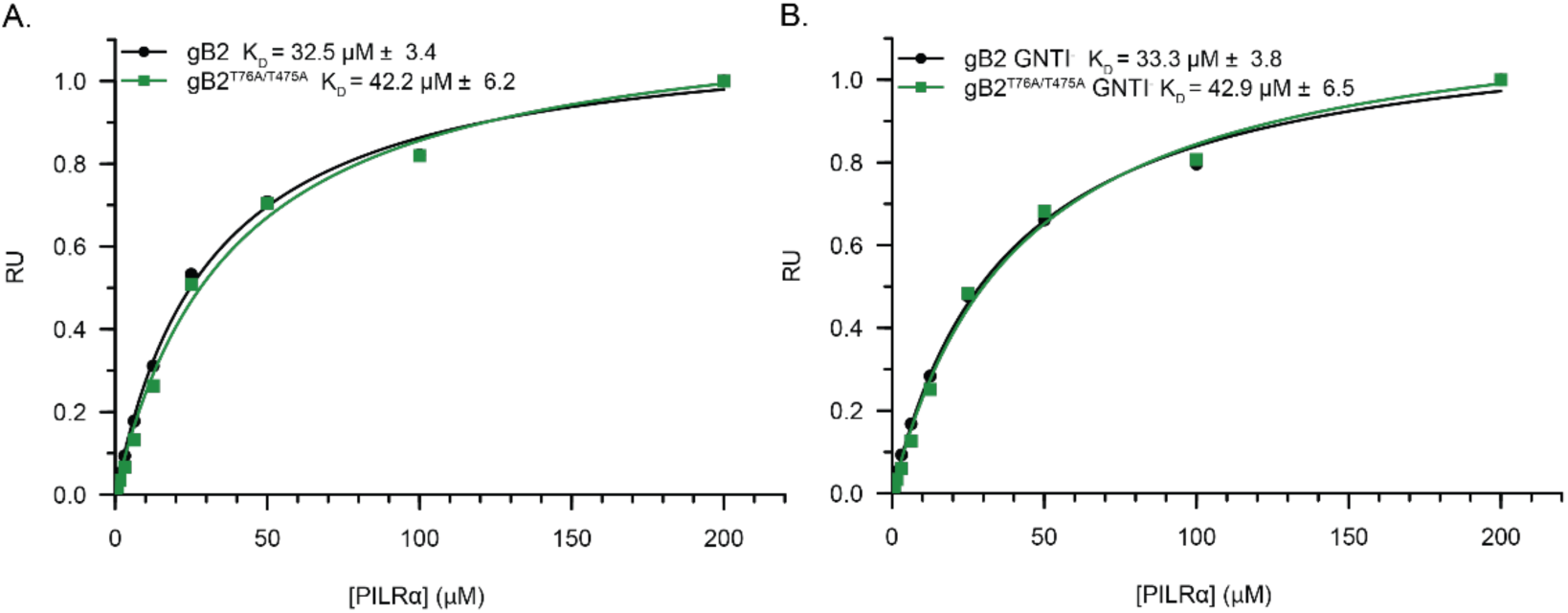
Multiple gB2 O-linked glycans play a role in PILRα interaction. SPR equilibrium datapoints, normalized to maximum binding, fitted to a 1:1 Langmuir binding model. A. Mutation of gB2 T76A/T475A slightly weakens PILRα binding. B. Interaction is unacected of PILRα binding to gB2 containing sialylated O-linked glycans but lacking sialylated N322 linked glycans.

### NMHCII-A binds to gB1 and gB2, but not gBV

NMHCII-A is notably different in structural and biochemical properties compared to MAG and PILRα. We determined a substantially higher affinity for the association of gB1 and gB2 with the rod-domain of NMHCII-A (K_D_ 0.627 ± 0.047 µM, 0.440 ± 0.030 µM) (Fig. 4G-H, Table 1 and Suppl. Fig. S3A-B). gBV does not interact with NMHCII-A (Suppl. Fig. S3C). The SPR data of NMHCII-A was fitted to a 1:1 kinetic model as equilibrium was not reached after each injection, precluding robust equilibrium-based analysis. gB2 expressed by GnTI−/− cells, containing high-mannose N-linked glycans, interacts with similar affinity and kinetics to NMHCII-A compared to gB2 containing a mix of N-linked glycans (Suppl. Fig. S4). The slow kinetics of gB1 and gB2 interaction with NMHCII-A, compared to the fast kinetics of MAG and PILRα and the much higher affinity suggest that the binding mode of gB1 and gB2 with NMHCII-A relies on protein-protein mediated interactions.

## Discussion

The three neurotropic human alphaherpesviruses HSV-1, HSV-2 and VZV are versatile viruses that can invade many cell types due to their adaptable interactions with host-cell receptors. Their interaction with the central nervous system is of particular interest in the context of severe neurological diseases such as HSE, Alzheimer’s Disease and Parkinson’s Disease. In this work, we probe interactions between essential fusogen glycoprotein B and host co-receptors MAG, PILRα and NMHC-IIA. Whereas the three gB variants share high sequence homology (ca. 90% between gB1-gB2, ca. 50% between gB1/2-gBV), variations in their PTMs as well as local protein motifs causes a specific host-virus interaction pattern that may determine individual cell tropism and neuroinflammation. Previous glycoproteomics studies of gB1 and gBV from authentic virions have mapped out the sites of O-linked glycosylation on gB in great detail and agree on the global distribution of sites, being concentrated in the flexible N-terminus and the flexible loop in domain II^49–51^. These studies used glycopeptide enrichment strategies that enables deeper coverage of low-occupancy sites spanning additional gB domains, but the lectins used for this enrichment required neuraminidase treatment to remove all sialic acids. The present work complements these studies by studying O-glycosylation with intact sialylation, providing additional information to survey candidate binding sites for MAG and PILRα to interpret the relative strength and specificity of their interactions.

To allow comparison between interaction affinities measured by SPR, we have minimized avidity enhanced interactions by employing the receptors, MAG, PILRα and NMHC-IIA as analyte and coupling the trimeric gB proteins to the sensor surface. From a physiological perspective, avidity will play an important role in the interaction of a herpesvirus with the receptors. We find that within a single gB chain the presence of multiple, well-spaced interactions sites (Fig. 3) would allow multiple MAG or PILRα receptors to bind. The interaction sites are exposed in both the pre– and post-fusion state of the gB proteins (Fig. 3). Trimerization of gB’s and the presence of a large number of gB trimers on the viral surface adds further avidity to increase the interaction strength^52^. By how much the relatively weak interactions that we determined for MAG and PILRα are enhanced in a cellular setting depends on the number of receptor molecules available for interaction with a single virus. MAG is the fifth highest expressed protein in myelin, with relative abundances determined between 1-4%^53^, indicating avidity enhanced interaction will likely play an important role.

All three gB variants we tested show similar, low micromolar affinity towards entry receptor MAG. MAG is known to preferentially bind α2,3-linked, and to a lesser extent α2,6 linked sialic acids^21,22^. Its ganglioside ligand GT1b possesses a NeuAcα(2-3)Galβ(1-3)GalNac branch, that is recognized by MAG^54^. MAG additionally binds oligosaccharides representing O-linked glycans that include α2,3 and/or α2,6 linked sialic acids, which outcompete N-linked glycans that contain α2,3 and/or α2,6 linked sialic acids, especially when the former are disialylated^22^. All gB variants contain monosialylated O-linked glycans (Fig. 2) and we observe monosialylated and disialylated N-linked glycans (Fig. 1). While we cannot distinguish the type of linkage in the glycoproteomics data, nor in the structural data due to flexibility of these glycan regions, it is likely that the MAG-gB interaction is primarily mediated by its O-glycans. Indeed, we find that gB2 lacking sialylated N-linked glycans, i.e. containing N-linked high-mannose glycans, bind with similar affinity to MAG (Fig. 5). All gB variants have multiple O-glycosylation sites, of which the most prominently sialylated sites are identified at the N-terminus and in domain II. The number and distribution of sialylated sites, in particular for gB2 (Fig. 2), most likely contribute to enhancing the affinity by an avidity effect for cell-surface expressed MAG. The relative contributions of the two binding locations with regards to functional viral entry remain elusive.

In contrast to MAG, PILRα bound only to gB1 and gB2, but not to gBV. Although the affinity of PILRα interaction with gB1 and gB2 is similar to the affinity of MAG with gB1 and gB2 (Fig. 4, Table 1), the specificity of this interaction is determined by O-glycan composition and likely by local protein sequence as well. Structural analysis of PILRα revealed additional contacts with a +1 proline residue relative to the O-linked glycosylation site^29,38,55^, which would narrow its binding specificity for gB over that of MAG. Previous work has shown that in HSV-1, PILRα-dependent infection enhancement is abrogated when two major O-glycan sites, T53 at the N-terminus and T480 in domain II, are mutated^36,37^. In our gB1 preparation we do not observe O-linked glycosylation at T53, while we do find sialylated glycosylation of T480. This suggests that the O-linked glycan on T480 is sufficient for binding PILRα in our assay. In contrast, we find three candidate PILRα binding sites in gB2, of which sialylated glycan occupancy at T76 and T475 (homologous to T480 in gB1) appear to be especially extensive. Mutation of gB2 T76 and T475 to alanines, however, only reduced affinity slightly (Fig. 6). The sustained binding of gB2^T76A/T475A^ to PILRα suggests the existence of an additional binding site for gB2. Indeed, gB2 has a serine-linked disialylated O-glycan followed by a proline at S55, which may contribute to PILRα binding, especially in absence of the T76 and T475 sites.

The role of PILRα in HSV-2 entry has been reported to be of little importance, while it is important for HSV-1^11,12^. Surprisingly, we find both gB’s interact with similar strength to PILRα and we identify multiple possible binding sites for PILRα on gB2. This suggests additional receptor(s) or regulatory mechanisms, such as heparan sulfate binding, are required for HSV entry. In contrast to HSV-1, for HSV-2 heparan sulfate attachment is mediated only by gB and not by gC^56^, pointing towards a role for glycosaminoglycan binding in HSV-2 entry. Studies regarding PILRα mediated entry always include heparan sulfate on host cells^11,13,14,57,58^. In gB2, two of the putative PILRα binding sites, T76 and S55, are flanking an N-terminal positively charged segment that has been suggested to be required for interaction with negatively charged glycosaminoglycans such as heparan sulfate, whereas for gB1, this positively charged segment is more distant to the putative PILRα binding sites. Possibly, glycosaminoglycans impair binding of HSV-2 to PILRα, explaining the lack of a role for PILRα in HSV-2 entry. In addition, the –2-asparagine relative to T475 in gB2 is N-linked glycosylated which may sterically hinder access of PILRα to this binding site, but not to T480 in gB1, that has no N-linked glycan nearby (Fig. 2). Whether PILRα plays a role in HSV-2 entry in absence of glycosaminoglycans remains to be determined.

NMHC-IIA binds to both gB1 and gB2, but not gBV. The affinity of the C-terminal coiled coil of NMHC-IIA to gB1 and gB2 is substantially higher compared to MAG and PILRα with an estimated K_D_ of 0.63 ± 0.05 µM and 0.44 ± 0.03 µM, respectively. The higher affinity interaction and the slower on– and off-rate for the binding kinetics suggest that binding is protein-protein mediated. In addition, there are no indications that NMHC-IIA can interact with glycans. It is currently not clear where the NMHC-IIA binding site on gB1 and gB2 is located but earlier work by others may give some leads. It was found that domain IV of gB1 was involved in receptor binding on heparan sulfate deficient CHO-cells^59,60^. Proteomics of these cells show expression of MAG and NMHC-IIA/B, but not PILRα^61^. Since MAG-gB interaction is dependent on glycans located largely in different areas of gB, antibody blocking of domain IV might prevent binding to NMHC-IIA/B and suggests domain IV of gB1/2 has a role in NMHC-IIA binding. This would be consistent with the observation that for VZV, for which we find gBV does not interact with NMHC-IIA, an antibody targeting domain IV but not the N-terminal domain did not affect VZV infection in MeWo cells^62^. Clearly, more work is required to pinpoint the NMHC-IIA binding site and detail the gB-NMHCIIA interaction mechanism.

To summarize, we probed the interactions between HSV-1, HSV-2 and VZV gB and their associated receptors MAG, PILRα and NMHC-IIA. Low micromolar affinity of gB variants to MAG and PILRα is characteristic for glycan-protein interactions. For MAG binding the α2,3-linked disialylated O-glycans located at the disordered N-terminus and domain II play an important role, although the sialylated N-linked glycans clustered at these sites may contribute as well. Binding of PILRα to gB1 and gB2 is explained by the presence of α2,6-linked sialylated O-glycans and +1 proline motif at the flexible N-terminus and flexible domain II loop. gBV lacks the combination of sialylated O-glycans with the +1 proline motif, which explains why PILRα does not bind to gBV. PILRα binding sites for gB1 and gB2 are of similar affinity but differ in their number and availability, possibly affected by the presence of an adjacent N-glycan site in domain II of gB2. The interaction of gB1 and gB2 with NMHC-IIA is substantially different, with stronger, submicromolar affinity and slower on and off rates. Precise mapping of these interactions may further enhance the understanding of herpesviruses utilizing these receptors and provide a basis for future therapeutic developments.

## Data availability

The raw LC-MS/MS files and analyses have been deposited to the ProteomeXchange Consortium via the PRIDE partner repository with the dataset identifier PXD073657. Cryo-EM density maps for gB2 have been deposited to the EMDB with identifier EMD-56537 and the structural model has been deposited to the PDB with 28IN.

## Acknowledgements

This work benefited from access to the Netherlands Centre for Electron Nanoscopy (NeCEN) at Leiden University. This research was funded by the Dutch Research Council NWO Gravitation 2013 BOO, Institute for Chemical Immunology (ICI; 024.002.009), as well as EM facility access through NEMI (184.034.014).

## Materials G Methods

### Construct design of expression plasmids

DNA sequences of HSV-1 gB (gB1, strain KOS, UNIPROT-ID P06437), HSV-2 gB (gB2, strain 333, UniProt ID P06763) and VZV gB (gBV, strain Dumas, Uniprot ID P09257) were ordered as synthetic DNA strings codon optimized for mammalian cell expression. PCR-amplification with 3’-NheI and 5’-BshTI restriction sites included in primers was performed to subclone ectodomain gB (residues 26-727 for gB1, 31-730 for gB2, 72-781 for gBV) DNA into mammalian cell expression vector prK5 (human IgG heavy chain signal peptide, C-terminal FOLDON trimerization domain followed by Strep-II-His_6_-tag). For gB2, fusion loop mutations A256S/F257S were made to increase solubility. Constructs for Surface Plasmon Resonance (SPR) immobilization were generated by PCR amplification with BamHI/NotI restriction sites to subclone into pUPE107.62 (cystatin secretion signal peptide, C-terminal biotin acceptor peptide (BAP) tag followed by His_6_-tag). Cloning of MAG (*M. Musculus*, Uniprot-ID P20917) ectodomain (residues 20-508) into mammalian cell expression vector pUPE107.03 (cystatin secretion signal peptide, C-terminal His_6_-tag) from IpA (Immunoprecise Antibodies) was described earlier^23^. Human PILRα (UniProt ID Q9UKJ1) and human NMHC-IIA (UniProt ID P35579) sequences were generated from Addgene plasmids 156577^63^ and 183512^64^ respectively. PILRα ectodomain (residues 20-154 and NMHC-IIA (residues 837-1960, i.e. the coiled-coil rod domain) DNA were generated by PCR with 3’ BamHI and 5’ NotI restriction sites to subclone into bacterial cell expression vector pETX13 (C-terminal His_6_-tag) or pETX30 (N-terminal His_6_-eGFPm tag) from IpA.

### Protein expression and purification of gB and MAG

HSV-1 gB, HSV-2 gB, VZV gB and MAG were expressed in Epstein-Barr virus nuclear antigen I (EBNA1)-expressing HEK293 cells. Recombinant gB protein with high-mannose N-glycans were expressed in N-acetylglucoaminyltransferase I-deficient (GnTI^−^) HEK293 cells (IpA). After 6 days, medium was harvested by spinning down at 5000xg for 20 minutes and filtered through a 0.22uM filter. For small scale purifications (4mL), medium was incubated with suspension Ni^2+^-sepharose excel beads (GE Healthcare) for 1-2 hours before batch purification by spinning down at 500xg for 5 minutes and subsequently washing with 2×10 bead volumes (BV) wash buffer (25mM Hepes pH 7.5, 500mM NaCl), 1×10 BV wash buffer (25 mM Hepes pH 7.5, 500mM NaCl, 5mM Imidazole) and eluted with elution buffer (25mM Hepes pH 7.5, 500mM NaCl, 500mM Imidazole). For large scale purification (1L), medium was run over a 5mL HisTrap^TM^ excel column (Cytiva) on an Äkta system sample pump before washing with >100mL wash buffer. Protein was eluted with elution buffer, fractions containing protein were pooled and concentrated to <500uL with an Amicon® ultra centrifugal filter (Merck) with a 100kDa cut off. Concentrated protein was run on a Superdex200 Increase column equilibrated with Size Exclusion Chromatography (SEC) buffer (25mM Hepes pH 7.5, 150mM NaCl) on an Äkta system. Protein containing fractions were pooled and concentrated to 1 mg/mL (gB) or 10 mg/mL (MAG), aliquoted and flash frozen to store at –80 °C. Details on protein expression and purification results can be found in Suppl. Fig. S6A-B for gB and Suppl. Fig. S7F-G for MAG.

### Protein expression and purification of PILRα and NMHC-IIA

Protein plasmids were transformed into E.coli BL21 competent expression cells (New England Biolabs) (NMHC-IIA) or 1:1 co-transformed with pGro7 plasmid expressing groES/groEL chaperone proteins (Takara Bio) (PILRα) and a single colony was picked for overnight inoculation in 50mL Lysogeny broth (LB) medium supplemented with 50 µg/mL kanamycin (NMHCII-A) or additionally, 20 µg/mL chloramphenicol (PILRα). 2-4 L cultures were inoculated to a starting OD280 of 0.05 in LB medium supplemented with 0.1% glucose to grow until OD280 0.6-0.8 for induction with 1mM (PILRα) or 0.5mM (NMHC-IIA) Isopropyl β-D-1-thiogalactopyranoside (IPTG). For PILRα, groES/groEL expression was induced with 1 mg/mL L-arabinose at an OD280 of 0.2. Cultures were left for expression at 25 °C (PILRα) or 18 °C (NMHC-IIA) overnight. Cells were harvested by centrifugation at 5000xg for 20 minutes at 4 °C, washed with PBS and cell pellets were kept at –20 °C until purification. To purify protein, cell pellets were thawed and resuspended in 10-20mL per liter of culture lysis buffer (PBS containing 0.1% Triton X-100, 1x Roche EDTA-free protease inhibitor cocktail tablets (Merck) and 10 µg/mL DNAse). For NMHC-IIA, lysis buffer and all subsequent buffers were supplemented with 1mM TCEP. Cells were lysed by sonification at 40% amplitude with 30s on, 30s off for 5 minutes. Lysate was cleared by first centrifuging at 5000xg for 20 minutes at 4 °C and a second centrifugation at 14000xg for 50 minutes at 4 °C before filtering through a 0.22 µm filter. Lysate was loaded on a 5 mL HisTrap^TM^ excel column (Cytiva) on an Äkta system sample pump and washed with >100mL wash buffer (25 mM Hepes pH 7.5, 500 mM NaCl). Elution was done with elution buffer (25 mM Hepes pH 7.5, 250 mM Imidazole). Protein containing fractions were pooled and concentrated using a 5 kDa cut off Amicon® ultra centrifugal filter (Merck) for PILRα or with a 100 kDa cut off membrane for NMHC-IIA to a volume <500 uL to load on a superdex75 Increase (PILRα) or a superose 6 Increase column (NMHC-IIA) run on an Äkta system equilibrated with SEC buffer (25 mM Hepes pH 7.5, 150 mM NaCl) or wash buffer for NMHC-IIA. Protein containing fractions were monitored with UV280 absorbance and UV488 if GFP is present and pooled and concentrated with the same respective filters. NMHC-IIA concentration was measured by performing a BCA assay following the instruction manual of a Pierce™ BCA Protein Assay kit (ThermoFischer Scientific). Details on protein expression and purification results can be found in Suppl. Fig. S7A-C for NMHC-IIA and Suppl. Fig. S7D-E for PILRα.

### Surface Plasmon Resonance (SPR)

HSV-1 gB, VZV gB, HSV-2 gB and HSV-2 gB mutant constructs were subcloned into mammalian expression vector pUPE107.62 containing a C-terminal biotin acceptor peptide (BAP) tag and His_6_-tag and co-transfected into EBNA1 expressing HEK293 cells or GnTI^−^ HEK293 cells 20:1 DNA ratio with E.coli biotin ligase BirA (sub-optimal secretion signal peptide, pUPE5.02) for *in vivo* biotinylation. A few hours after transfection, additional sterile biotin (3 mg/mL in Tris-HCl buffer) was added to the culture (30 µL per 4 mL culture). Purification of ligand constructs was done by performing small scale Ni-excel affinity purification (see Suppl. Fig. S6C for results) and diluted to a concentration series of 400, 200 and 100 nM before spotting on a P-STREP SensEye® (Ssens) chip using a Continuous Flow Microspotter (Wasatch Microfliudics). For neuraminidase treatment, imidazole was removed using Zeba™ Spin Desalting Columns (ThermoFischer Scientific) before addition of Glycobuffer 1 (New England Biolabs) to match 1x buffer concentration and 2 µL α2,3,6,8-Neuraminidase (100 units, New England Biolabs) per 1 µg protein, after which samples were left at room temperature overnight. Buffer used for dilutions and for SPR measurements consisted of 25 mM Hepes pH 7.5, 150 mM NaCl and 0.005% Tween-20 or, in the case of NMHCII-A, 25 mM Hepes pH 7.5, 500 mM NaCl and 0.005% Tween-20. After spotting, the surface of the chip was quenched with 1 mg/mL biotin. SPR measurements were performed on a MX96 SPRi instrument (IBIS Technologies) with a constant temperature of 25 °C. Three spots with different ligand concentrations (200, 100, 50 nM) were used as technical replicates for data processing. Data were processed with SPRINTX software (IBIS Technologies) for referencing and baseline correction and Scrubber (BioLogic Software) for equilibrium sensogram processing. Equilibrium data values were fitted to a 1:1 Langmuir binding curve using non-linear regression in Rstudio according to the following equation:

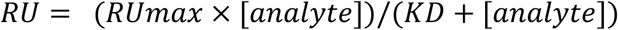

Where RUmax is the maximum analyte binding. Standard deviations for each K_D_ calculation were produced by Rstudio. For kinetic affinity curves, SPR sensograms were fitted to a 1:1 binding model in Scrubber. Standard deviation values were provided by Scrubber. Mean standard deviations were calculated as square root of the sum of the squared standard deviation values divided by the total number of standard deviations.

### Glycoproteomics

5 µg purified protein was used for each digestion replicate. Protein was diluted in digestion buffer (1% SDC, 100 mM Tris-HCl, 10 mM TCEP, 40 mM CAA, pH 8) and incubated at 95 °C for 20 minutes prior to RT incubation in the dark for 25 minutes. For O-glycoproteomics, samples were diluted 2x in PBS containing 4 µg PNGaseF and incubated at 37 °C overnight. Subsequently, protein samples were diluted 1:5 in 50 mM AMBIC and respective digestion enzymes (trypsin, GluC + trypsin, aphalytic protease (aLP) or chymotrypsin) were added in a 1:50 enzyme:protein w/w ratio. Protein was digested overnight at 37 °C and acidified by adding TFA to a final concentration of 1% and centrifuged at 21.000xg for 20 minutes at RT. Supernatant was desalted using an Oasis HLB desalting plate (Waters) prepared with 2x 100 µL 100% acetonitrile (ACN) and 2x 100 µL 0.1% TFA and after loading supernatant, column material was washed by adding 2x 100 µL 0.1% TFA and protein peptides were eluted by addition of 2x 50 µL 50% ACN/0.1% TFA. Collected samples were lyophilized in a SpeedVac vacuum concentrator (ThermoFischer Scientific) and kept at –20 °C for storage.

Peptides were analyzed on a nanospray UHPLC system Ultimate 3000 (ThermoFisher Scientific) coupled online to an Orbitrap Eclipse Mass Spectrometer (Thermo Fisher Scientific). Peptides were loaded onto an Acclaim Pepmap 100 C18 (5 mm × 0.3 mm, 5 μm, ThermoFisher Scientific) trap column at 32 °C and separated on an in-house made analytical column with integrated emitter (ReproSil-Pur 120 C18, 50 cm × 75 μm, 1.9 μm, Dr. Maisch) at 50 °C. The LC mobile phase A was 0.1% formic acid and B 0.1% formic acid in 80% acetonitrile. The flowrate was 300 nL/min on a 44 min gradient (4% – 55% B). MS analysis was performed using HCD, HCD-pd-sHCD and HCD-pd-EThcD methods. MS1 settings were as follows: 120,000 resolution; 350-2000 m/z scan range; 100% normalized AGC target; 246 ms maximum injection time. MS2 settings were as follows: 30,000 resolution; 120-4000 m/z scan range; 50 ms maximum injection time; 100% normalized AGC target; 29% NCE. For the HCD-pd-sHCD and HCD-pd-EThcD methods, additional sHCD or EThcD scans were acquired when the HCD contained at least 3 oxonium ion masses, as specified in Table 2. The sHCD and EThcD settings were as follows: 60,000 resolution; 120-4000 m/z scan range; 150 ms maximum injection time; 100% normalized AGC target. In sHCD, NCE was set to stepped mode, using 10, 25 and 40% NCE. In EThcD, calibrated ETD settings were used and NCE of 15%.

**Table 2.** Oxonium ions to trigger additional MS2. Abbreviation of glycan compositions: Hex: hexose, NAc: N-acetyl hexosamine, P: phosphomannose, dHex: deoxyhexose, NeuAc: N-acetylneuraminic acid.

| Compound name | m/z |
| --- | --- |
| Hex | 127.039 |
| Hex | 145.0495 |
| Hex | 163.0601 |
| PHex | 243.0264 |
| Phex2 | 405.0793 |
| HexNAc | 138.055 |
| HexNAc | 168.0655 |
| HexNAc | 186.0761 |
| HexNAc | 204.0847 |
| NeuAc | 274.0921 |
| NeuAc | 292.1027 |
| HexHexNAc | 366.1395 |
| HexNAc2 | 407.166 |
| HexHexNAcdHex | 512.1974 |
| NeuAcHexHexNAc | 657.2349 |

The acquired data was analyzed using Byonic (v5.9.125) against a custom database of recombinant gB sequences and the proteases used in the experiment, searching for glycan modifications with 10/20 ppm search windows for MS1/MS2, respectively. Up to 3 missed cleavages were permitted using C-terminal cleavage at R/K for trypsin, 5 at R/K/E/D for trypsin-gluC, and 5 at WFLYM for chymotrypsin. For N-linked analysis, carbamidomethylation of cysteine was set as fixed modification, oxidation of methionine/tryptophan as variable rare 2. N-glycan modifications were set as variable common 2, allowing up to max. 2 variable common and 2 rare modifications per peptide. All N-linked glycan databases from Byonic were merged into a single non-redundant list to be included in the database search. The Byonic result files were filtered for glycopeptides with scores ≥ 200, and additional peptide-spectrum matches (PSMs) containing non-glycosylated asparagines were included at the same score cutoff. Glycans were classified based on HexNAc content as truncated (≤ 2 HexNAc; < 3 Hex), paucimannose (2 HexNAc, 3 Hex), high mannose (2 HexNAc; > 3 Hex), hybrid (3 HexNAc) or complex (> 3 HexNAc). For O-linked glycoproteomics analyses, settings were as specified above, with the additional inclusion of deamidation of asparagine as variable modification, and replacing the N-linked glycan database with a list of 17 common O-linked glycan modifications set as ‘variable common 3’. The deposition including all raw files, Byonic results, databases and scripts use for glycopeptide counting and classification can be accessed under PXD073657 with token 80QiYirnruNw.

### Cryo-EM sample preparation

Purified HSV-2 glycoprotein B was diluted in SEC buffer (25 mM Hepes pH 7.5, 150 mM NaCl) to 0.3 mg/mL and plunge frozen using a Vitrobot Mark IV (ThermoFischer Scientific) set to 100% humidity at 8 °C. Sample was applied to a glow discharged (30 seconds at 15 mA) 300 mesh R1.2/R1.3 copper grid (QuantiFoil) before blotting for 4.5 seconds with blot force 0. Grids were rapidly frozen in a liquid ethane/propane mixture. Data collection was done at a 300 kV Titan Krios Electron Microscope (ThermoFischer Scientific) equipped with a K3 direct electron summit camera (Gatan). 4611 movies were collected at a magnification of 105.000x with 50 frames and a total dose of 50 e^−^/Å^2^ in superresolution mode corresponding to a pixel size of 0.418 Å, with a defocus range of –2.3 to –1.1 µM in steps of 0.3 µm.

### Cryo-EM data processing

Data were processed with C3 symmetry using CryoSparc v3.3.1 to v4.5.3 software^65–67^. Movies were processed using Patch Motion Correction and binned 2x to a physical pixel size of 0.836. After Patch CTF correction, movies were manually curated leaving 3846 corrected micrographs (765 rejected). To generate templates, particles were picked on 50 micrographs using Blob picker with an elliptical blob and minimum particle diameter of 60 Å and maximum particle diameter 220 Å. After inspection of picks, 951 of 18.009 picks were included (NCC score > 0,480, local power between 310,000 and 589,000) and extracted using a box size of 410 pixels downsized to 256 after which 5702 particles remained. The particles were 2D classified to 50 classes of which 14 classes were selected for template picking on all micrographs using a particle diameter of 200 Å. After inspection of picks (NCC score > 0,490, local power between 910,000 and 1428,000), 276.382 particles were extracted using a box size of 410 pixels downsized to 256 and subjected to 2D classification. 211.067 particles were used for Ab-initio reconstruction of 3 volumes, which were used for 2 rounds of heterogeneous refinement at C3 symmetry using the initial 276.382 extracted particles. 147.888 good particles were used for Non-Uniform (NU) Refinement yielding a 2.9 Å resolution structure. Particles were subjected to local CTF refinement and re-extracted to a box size of 410 and subsequently used for ab initio reconstruction and NU refinement resulting in a final 2.78 Å resolution volume. Volumes were assessed and figures were made using ChimeraX v.1.8^68–70^.

### Refinement of the HSV-2 gB model

A homology model of HSV-2 gB based on postfusion HSV-1 gB (PDB ID: 2GUM) was generated by SWISS-MODEL^71^ and used for refinement with PHENIX^72^ in the CCP-EM^73,74^ environment. Model was manually checked, glycan moieties were added using COOT ^75^. Glycan conformations were validated using the Privateer web app^76^.

## Notes

### Competing Interest Statement

The authors have declared no competing interest.

## References

1. Gnann, J. W. & Whitley, R. J. Herpes Simplex Encephalitis: an Update. Current Infectious Disease Reports vol. 19 Preprint at 10.1007/s11908-017-0568-7 (2017).

2. Marcocci, M. E. et al. Herpes Simplex Virus-1 in the Brain: The Dark Side of a Sneaky Infection. Trends in Microbiology vol. 28 808–820 Preprint at 10.1016/j.tim.2020.03.003 (2020).

3. Eyting, M. et al. A natural experiment on the effect of herpes zoster vaccination on dementia. Nature 641, 438–446 (2025).

4. Bjornevik, K., Münz, C., Cohen, J. I. & Ascherio, A. Epstein–Barr virus as a leading cause of multiple sclerosis: mechanisms and implications. Nature Reviews Neurology vol. 19 160–171 Preprint at 10.1038/s41582-023-00775-5 (2023).

5. Lanz, T. V. et al. Clonally expanded B cells in multiple sclerosis bind EBV EBNA1 and GlialCAM. Nature 603, 321–327 (2022).

6. Steel, A. J. & Eslick, G. D. Herpes viruses increase the risk of Alzheimer’s disease: A meta-analysis. Journal of Alzheimer’s Disease 47, 351–364 (2015).

7. Akhtar, J. & Shukla, D. Viral entry mechanisms: Cellular and viral mediators of herpes simplex virus entry. FEBS Journal vol. 276 7228–7236 Preprint at 10.1111/j.1742-4658.2009.07402.x (2009).

8. Cheshenko, N. & Herold, B. C. Glycoprotein B Plays a Predominant Role in Mediating Herpes Simplex Virus Type 2 Attachment and Is Required for Entry and Cell-to-Cell Spread. Journal of General Virology vol. 83 (2002).

9. Suenaga, T. et al. Myelin-associated glycoprotein mediates membrane fusion and entry of neurotropic herpesviruses. Proc. Natl. Acad. Sci. U. S. A. 107, 866–871 (2010).

10. Cole, N. L. & Grose, C. Membrane fusion mediated by herpesvirus glycoproteins: The paradigm of varicella-zoster virus. Reviews in Medical Virology vol. 13 207–222 Preprint at 10.1002/rmv.377 (2003).

11. Arii, J. et al. Entry of Herpes Simplex Virus 1 and Other Alphaherpesviruses via the Paired Immunoglobulin-Like Type 2 Receptor α. J. Virol. 83, 4520– 4527 (2009).

12. Shukla, S. Y., Singh, Y. K. & Shukla, D. Role of nectin-1, HVEM, and PILR-in HSV-2 entry into human retinal pigment epithelial cells. Invest. Ophthalmol. Vis. Sci. 50, 2878–2887 (2009).

13. Satoh, T. et al. PILRα Is a Herpes Simplex Virus-1 Entry Coreceptor That Associates with Glycoprotein B. Cell 132, 935–944 (2008).

14. Fan, Q. & Longnecker, R. The Ig-Like V-Type Domain of Paired Ig-Like Type 2 Receptor Alpha Is Critical for Herpes Simplex Virus Type 1-Mediated Membrane Fusion. J. Virol. 84, 8664–8672 (2010).

15. Lathe, R. & Haas, J. G. Distribution of cellular HSV-1 receptor expression in human brain. J. Neurovirol. 23, 376–384 (2017).

16. Arii, J. et al. Non-muscle myosin IIA is a functional entry receptor for herpes simplex virus-1. Nature 467, 859–862 (2010).

17. Wang, C. et al. Nectin-1 and Non-muscle Myosin Heavy Chain-IIB: Major Mediators of Herpes Simplex Virus-1 Entry Into Corneal Nerves. Front. Microbiol. 13, (2022).

18. Arii, J., Hirohata, Y., Kato, A. & Kawaguchi, Y. Nonmuscle Myosin Heavy Chain IIB Mediates Herpes Simplex Virus 1 Entry. J. Virol. 89, 1879–1888 (2015).

19. Dai, Y. et al. TMEFF1 is a neuron-specific restriction factor for herpes simplex virus. Nature 632, 383–389 (2024).

20. Clarke, R. W. Forces and Structures of the Herpes Simplex Virus (HSV) Entry Mechanism. ACS Infect. Dis. 1, 403–415 (2016).

21. Kelm, S. et al. Sialoadhesin, Myelin-Associated Glycoprotein and CD22 Define a New Family of Sialic Acid-Dependent Adhesion Molecules of the Immunoglobulin Superfamily. Current Biology vol. 4 (1994).

22. Blixt, O., Collins, B. E., Van den Nieuwenhof, I. M., Crocker, P. R. & Paulson, J. C. Sialoside specificity of the siglec family assessed using novel multivalent probes: Identification of potent inhibitors of myelin-associated glycoprotein. Journal of Biological Chemistry 278, 31007–31019 (2003).

23. Pronker, M. F. et al. Structural basis of myelin-associated glycoprotein adhesion and signalling. Nat. Commun. 7, (2016).

24. David, S., Kottis, V., Dunn, R. & Braunt, P. E. Identification of Myelin-Associated Glycoprotein as a Major Myelin-Berived of Neurite Growth. Neuron vol. 13 (1994).

25. Fournier, N. et al. FDF03, a Novel Inhibitory Receptor of the Immunoglobulin Superfamily, Is Expressed by Human Dendritic and Myeloid Cells. The Journal of Immunology 165, 1197–1209 (2000).

26. Lee, K. J. et al. Paired Ig-like type 2 receptor-derived agonist ligands ameliorate inflammatory reactions by downregulating β1 integrin activity. Mol. Cells 39, 557–565 (2016).

27. Kohyama, M. et al. Monocyte infiltration into obese and fibrilized tissues is regulated by PILRα. Eur. J. Immunol. 46, 1214–1223 (2016).

28. Sun, Y. et al. Evolutionarily conserved paired immunoglobulin-like receptor α (PILRα) domain mediates its interaction with diverse sialylated ligands. Journal of Biological Chemistry 287, 15837–15850 (2012).

29. Kuroki, K. et al. Structural basis for simultaneous recognition of an O-glycan and its attached peptide of mucin family by immune receptor PILRα. Proc. Natl. Acad. Sci. U. S. A. 111, 8877–8882 (2014).

30. Furukawa, X. A. et al. Structural and thermodynamic analyses reveal critical features of glycopeptide recognition by the human PILRα immune cell receptor. Journal of Biological Chemistry 292, 21128–21136 (2017).

31. Xia, L. et al. PILRα on tumor cells interacts with the T cell surface protein CD99 to suppress antitumor immunity. *Nat*. Cancer 6, 1184–1201 (2025).

32. Javier-Torrent, M. & Saura, C. A. Conventional and Non-Conventional Roles of Non-Muscle Myosin II-Actin in Neuronal Development and Degeneration. Cells vol. 9 Preprint at 10.3390/cells9091926 (2020).

33. Vicente-Manzanares, M., Ma, X., Adelstein, R. S. & Horwitz, A. R. Non-muscle myosin II takes centre stage in cell adhesion and migration. Nature Reviews Molecular Cell Biology vol. 10 778–790 Preprint at 10.1038/nrm2786 (2009).

34. Phillips, C. L., Yamakawa, K. & Adelstein, R. S. Cloning of the CDNA Encoding Human Nonmuscle Myosin Heavy Chain-B and Analysis of Human Tissues with Isoform-Specific Antibodies. Journal of Muscle Research and Cell Motility vol. 16 (1995).

35. Suenaga, T. et al. Sialic acids on varicella-zoster virus glycoprotein B are required for cell-cell fusion. Journal of Biological Chemistry 290, 19833–19843 (2015).

36. Arii, J. et al. A Single-Amino-Acid Substitution in Herpes Simplex Virus 1 Envelope Glycoprotein B at a Site Required for Binding to the Paired Immunoglobulin-Like Type 2 Receptor α (PILRα) Abrogates PILRα-Dependent Viral Entry and Reduces Pathogenesis. J. Virol. 84, 10773–10783 (2010).

37. Wang, J. et al. Binding of Herpes Simplex Virus Glycoprotein B (gB) to Paired Immunoglobulin-Like Type 2 Receptor α Depends on Specific Sialylated O – Linked Glycans on gB. J. Virol. 83, 13042–13045 (2009).

38. Lu, Q., et al. PILRα and PILRβ have a siglec fold and provide the basis of binding to sialic acid. Proc. Natl. Acad. Sci. U. S. A. 111, 8221–8226 (2014).

39. Suenaga, T., Mori, Y., Suzutani, T. & Arase, H. Siglec-7 mediates varicella-zoster virus infection by associating with glycoprotein B. Biochem. Biophys. Res. Commun. 607, 67–72 (2022).

40. Vollmer, B., et al. The Prefusion Structure of Herpes Simplex Virus Glycoprotein B. Sci. Adv vol. 6 www.compbio.dundee.ac.uk/aacon/ (2020).

41. Hulswit, R. J. G. et al. Cetacean coronavirus spikes highlight S glycoprotein structural plasticity. Preprint at 10.1101/2025.05.20.655071 (2025).

42. Peng, W. et al. Glycan shield of the ebolavirus envelope glycoprotein GP. *Commun*. Biol. 5, (2022).

43. Chongsaritsinsuk, J. et al. Quantification and Site-Specific Analysis of Co-occupied N– and O-Glycopeptides. J. Proteome Res. 23, 5449–5461 (2024).

44. Tian, W. et al. O-glycosylation pattern of the SARS-CoV-2 spike protein reveals an “O-Follow-N” rule. Cell Research vol. 31 1123–1125 Preprint at 10.1038/s41422-021-00545-2 (2021).

45. Vollmer, B. et al. A nanobody specific to prefusion glycoprotein B neutralizes HSV-1 and HSV-2. Nature https://doi.org/10.1038/s41586-025-09438-5 (2025) doi:10.1038/s41586-025-09438-5.

46. Wang, G. et al. A broadly neutralizing antibody confers cross-genus protection against alphaherpesviruses by inhibiting gB-mediated membrane fusion. Nat. Commun. 16, 11144 (2025).

47. Seyfizadeh, N. et al. Development of a highly effective combination monoclonal antibody therapy against Herpes simplex virus. J. Biomed. Sci. 31, (2024).

48. Roark, R. S. et al. Prefusion structure, evasion and neutralization of HSV-1 glycoprotein B. Nat. Microbiol. 10, 2966–2980 (2025).

49. Bagdonaite, I. et al. Global mapping of o-glycosylation of varicella zoster virus, human cytomegalovirus, and Epstein-Barr virus. Journal of Biological Chemistry 291, 12014–12028 (2016).

50. Bagdonaite, I. et al. A Strategy for O-Glycoproteomics of Enveloped Viruses—the O-Glycoproteome of Herpes Simplex Virus Type 1. PLoS Pathog. 11, (2015).

51. Bagdonaite, I. et al. Glycoengineered keratinocyte library reveals essential functions of specific glycans for all stages of HSV-1 infection. Nat. Commun. 14, (2023).

52. Handler, C. G., Eisenberg, R. J. & Cohen, G. H. Oligomeric Structure of Glycoproteins in Herpes Simplex Virus Type 1. JOURNAL OF VIROLOGY vol. 70 https://journals.asm.org/journal/jvi (1996).

53. Jahn, O., Tenzer, S. & Werner, H. B. Myelin proteomics: Molecular anatomy of an insulating sheath. Molecular Neurobiology vol. 40 55–72 Preprint at 10.1007/s12035-009-8071-2 (2009).

54. Vinson, M. et al. Myelin-associated glycoprotein interacts with ganglioside GT1b. A mechanism for neurite outgrowth inhibition. Journal of Biological Chemistry 276, 20280–20285 (2001).

55. Agostini, S. et al. The PILRA G78R Variant Correlates with Higher HSV-1-Specific IgG Titers in Alzheimer’s Disease. Cell. Mol. Neurobiol. **39**, 1217– 1221 (2019).

56. Gerber, S. I., Belval, B. J. & Herold, B. C. Differences in the Role of Glycoprotein C of HSV-1 and HSV-2 in Viral Binding May Contribute to Serotype Differences in Cell Tropism. VIROLOGY vol. 214 (1995).

57. Shah, A., Farooq, A. V, Tiwari, V., Kim, M.-J. & Shukla, D. HSV-1 Infection of Human Corneal Epithelial Cells: Receptor-Mediated Entry and Trends of Re-Infection. http://www.molvis.org/molvis/v16/a265 (2010).

58. Chowdhury, S., Chouljenko, V. N., Naderi, M. & Kousoulas, K. G. The Amino Terminus of Herpes Simplex Virus 1 Glycoprotein K Is Required for Virion Entry via the Paired Immunoglobulin-Like Type-2 Receptor Alpha. J. Virol. 87, 3305–3313 (2013).

59. Atanasiu, D. et al. Bimolecular Complementation Defines Functional Regions of Herpes Simplex Virus gB That Are Involved with gH/gL as a Necessary Step Leading to Cell Fusion. J. Virol. 84, 3825–3834 (2010).

60. Bender, F. C., Whitbeck, J. C., Lou, H., Cohen, G. H. & Eisenberg, R. J. Herpes Simplex Virus Glycoprotein B Binds to Cell Surfaces Independently of Heparan Sulfate and Blocks Virus Entry. J. Virol. 79, 11588–11597 (2005).

61. Heffner, K. et al. Expanded Chinese hamster organ and cell line proteomics profiling reveals tissue-specific functionalities. Sci. Rep. 10, (2020).

62. Oliver, S. L. et al. The N-terminus of varicella-zoster virus glycoprotein B has a functional role in fusion. Preprint at 10.1101/2020.09.08.269191 (2020).

63. Wojtowicz, W. M. et al. A Human IgSF Cell-Surface Interactome Reveals a Complex Network of Protein-Protein Interactions. Cell 182, 1027–1043.e17 (2020).

64. Panza, E. et al. Transfection of the mutant MYH9 cDNA reproduces the most typical cellular phenotype of MYH9-related disease in different cell lines. Pathogenetics 1, (2008).

65. Punjani, A., Rubinstein, J. L., Fleet, D. J. & Brubaker, M. A. CryoSPARC: Algorithms for rapid unsupervised cryo-EM structure determination. Nat. Methods 14, 290–296 (2017).

66. Rubinstein, J. L. & Brubaker, M. A. Alignment of cryo-EM movies of individual particles by optimization of image translations. J. Struct. Biol. 192, 188–195 (2015).

67. Punjani, A., Zhang, H. & Fleet, D. J. Non-uniform refinement: adaptive regularization improves single-particle cryo-EM reconstruction. Nat. Methods 17, 1214–1221 (2020).

68. Meng, E. C. et al. UCSF ChimeraX: Tools for structure building and analysis. Protein Science 32, (2023).

69. Pettersen, E. F. et al. UCSF ChimeraX: Structure visualization for researchers, educators, and developers. Protein Science 30, 70–82 (2021).

70. Goddard, T. D. et al. UCSF ChimeraX: Meeting modern challenges in visualization and analysis. Protein Science 27, 14–25 (2018).

71. Waterhouse, A. et al. SWISS-MODEL: Homology modelling of protein structures and complexes. Nucleic Acids Res. 46, W296–W303 (2018).

72. Afonine, P. V. et al. Towards automated crystallographic structure refinement with phenix.refine. Acta Crystallogr. D Biol. Crystallogr. 68, 352– 367 (2012).

73. Burnley, T., Palmer, C. M. & Winn, M. Recent developments in the CCP-EM software suite. in Acta Crystallographica Section D: Structural Biology vol. 73 469–477 (International Union of Crystallography, 2017).

74. Wood, C. et al. Collaborative computational project for electron cryo-microscopy. Acta Crystallogr. D Biol. Crystallogr. 71, 123–126 (2015).

75. Emsley, P., Lohkamp, B., Scott, W. G. & Cowtan, K. Features and development of Coot. Acta Crystallogr. D Biol. Crystallogr. 66, 486–501 (2010).

76. Dialpuri, J. S. et al. Online carbohydrate 3D structure validation with the Privateer web app. Acta Crystallogr. F Struct. Biol. Commun. 80, 30–35 (2024).

