## Supplementary Information for "The glycan shield of alphaherpesvirus glycoprotein B modulates host co-receptor binding"

**Contents:**

- Legends for Supplementary Data S1-2
- Supplementary Table S1
- Supplementary Figures S1-7

**Supplementary Data S1.** Overview of N-glycoproteomics results for gB1, gB2, and gBV. Summaries list total filtered glycopeptide counts, categorized by HexNAc content, by absolute count and fraction of the total. Filtered spectra overviews show all included glycopeptides drawn from Byonic results.

**Supplementary Data S2.** Overview of O-glycoproteomics results for gB1, gB2, and gBV. Summaries list total filtered glycopeptide counts, summed by glycan composition. Filtered spectra overviews show all included glycopeptides drawn from Byonic results.

**Supplementary Table S1.** Data collection and refinement statistics for gB2 postfusion cryo-EM reconstruction and model.

| | | |
|---|---|---|
| **Data collection and processing** | | |
| Magnification | | 105kx |
| Voltage (kV) | | 300 |
| Electron exposure ($e^{-}$/ Å$^2$) | | 50 |
| Defocus range (μm) | | -2.3 to -1.1 |
| **Pixel size, super resolution (Å)** | | 0.418 |
| Symmetry imposed | | C3 |
| Map resolution (Å) | | 2.76 |
| FSC threshold | | 0.143 |
| **Refinement** | | |
| Initial model used | | SWISS-MODEL homology model based on PDB-ID: 2GUM |
| B-factors (Å$^2$) | | |
| | Protein | 92.4 |
| | Ligand | 132.7 |
| #non-hydrogen atoms | | 14748 |
| #residues | | 1806 |
| R.m.s. deviations | | |
| | Bond lengths (Å) | 0.004 |
| | Bond angels (degrees) | 0.91 |
| Validation | | |
| | Molprobity score | 1.59 |
| | Clash score | 6.11 |
| | Poor rotamers (%) | 1.92 |
| Ramachandran plot | | |
| | Favored (%) | 97.83 |
| | Allowed (%) | 1.84 |
| | Outliers (%) | 0.33 |

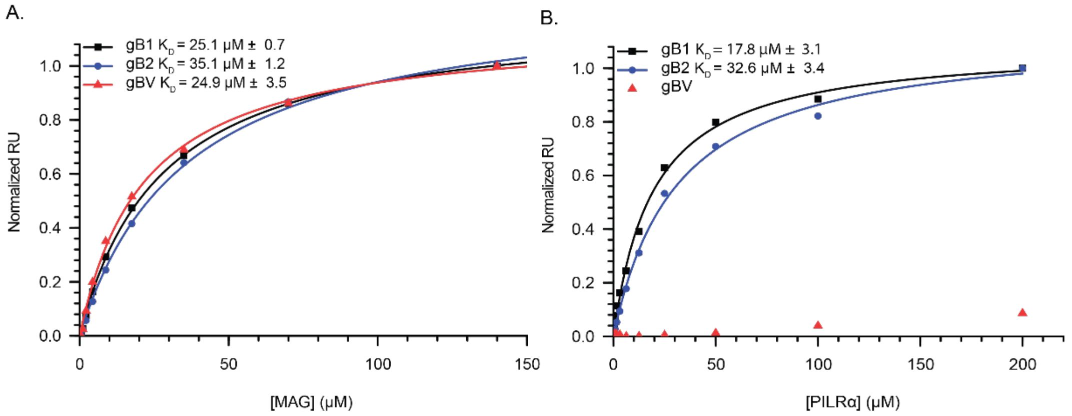


**Supplementary Figure S1.** Equilibrium binding affinity curves of receptor binding to gB1, gB2 and gBV. Data were normalized to their respective maximum RU value and fit to a 1:1 Langmuir binding curve to calculate $K_D$ values. A. MAG binds to gB1, gB2 and gBV with low μM affinity. B. PILRα binds to gB1 and gB2 with low μM affinity, but gBV does not bind to PILRα. Data for gBV was normalized to the maximum RU value of gB1 for visualization purposes.

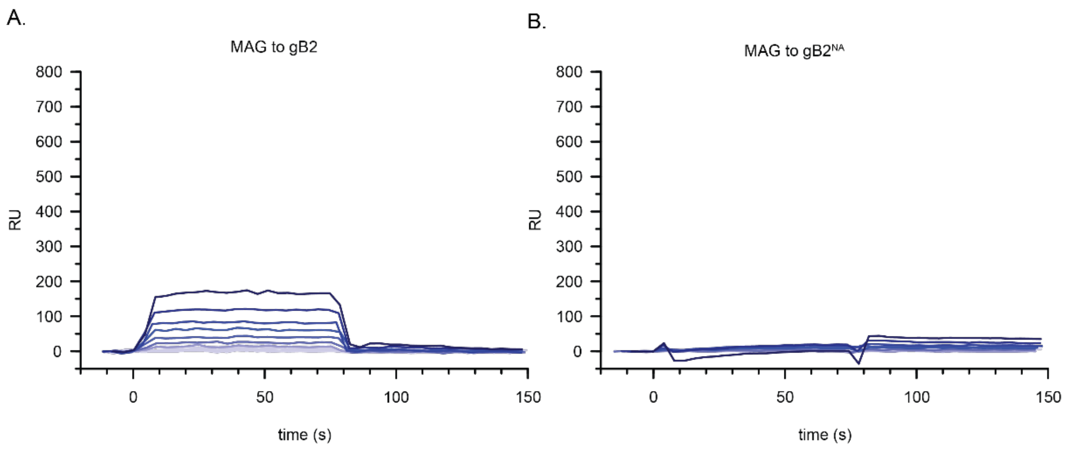


**Supplementary Figure S2.** SPR sensograms for MAG binding to gB2 at two-fold dilutions with a maximum concentration of 140 µM MAG. A. MAG binds to untreated gB2. B. MAG binding is abrogated after sufficient sialic acid trimming of gB2.

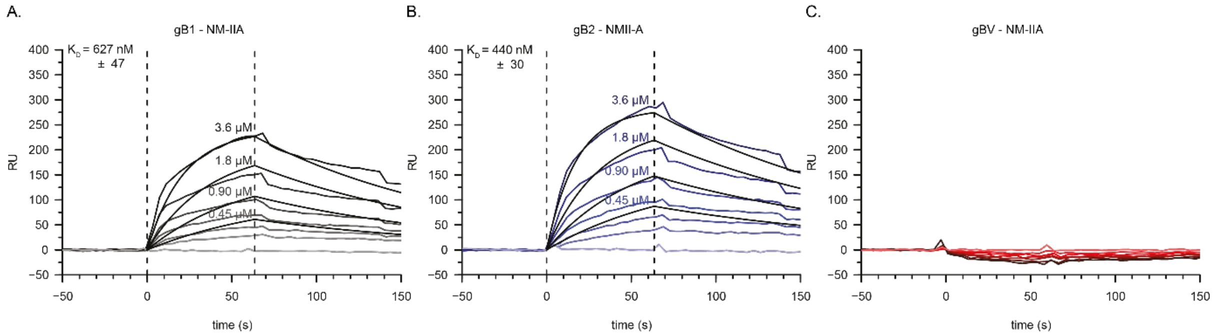


**Supplementary Figure S3.** SPR sensograms with fitted 1:1 kinetic curves of NM-IIA binding to gB variants. A. NM-IIA binds to gB1 with 627 nM affinity. B. NM-IIA binds to gB2 with 440 nM affinity. C. NM-IIA does not bind to gBV.

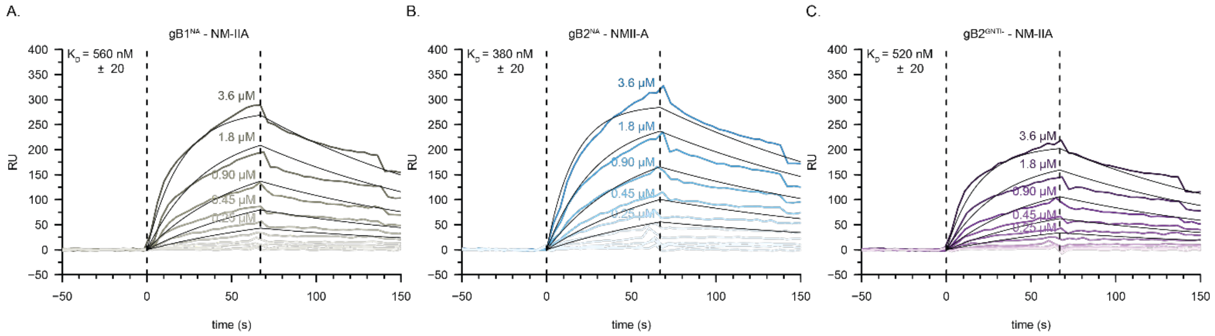


**Supplementary Figure S4.** Glycan composition of gB1 and gB2 does not influence binding capacity to NM-IIA. A. SPR sensograms with fitted 1:1 kinetic binding curves to gB1 treated with neuraminidase. B. SPR sensograms with fitted 1:1 kinetic binding curves to gB2 treated with neuraminidase. C. SPR sensograms with fitted 1:1 kinetic binding curves to gB2 produced in GNTI$^{-/-}$ cells, i.e. with high mannose N-glycans.

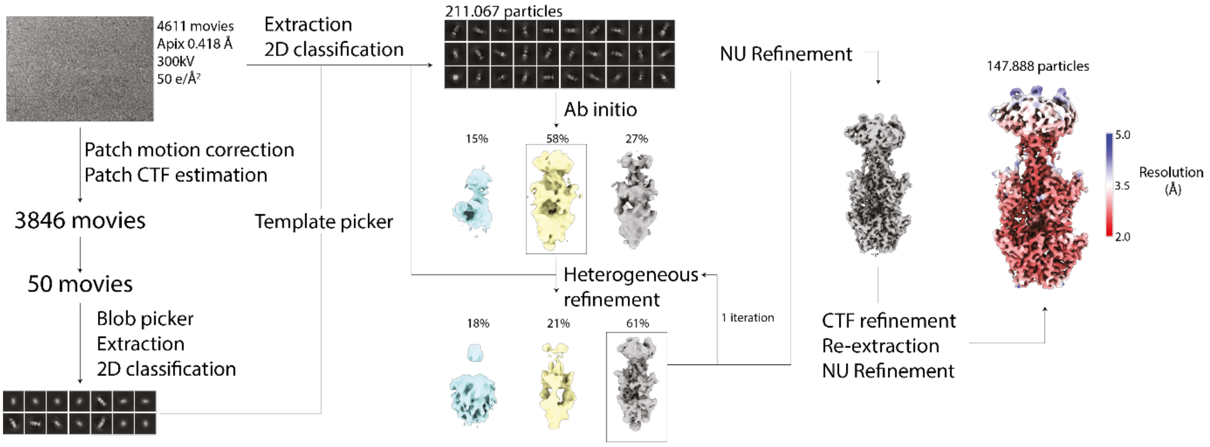


**Supplementary Figure S5.** Cryo-EM data processing scheme for gB2.

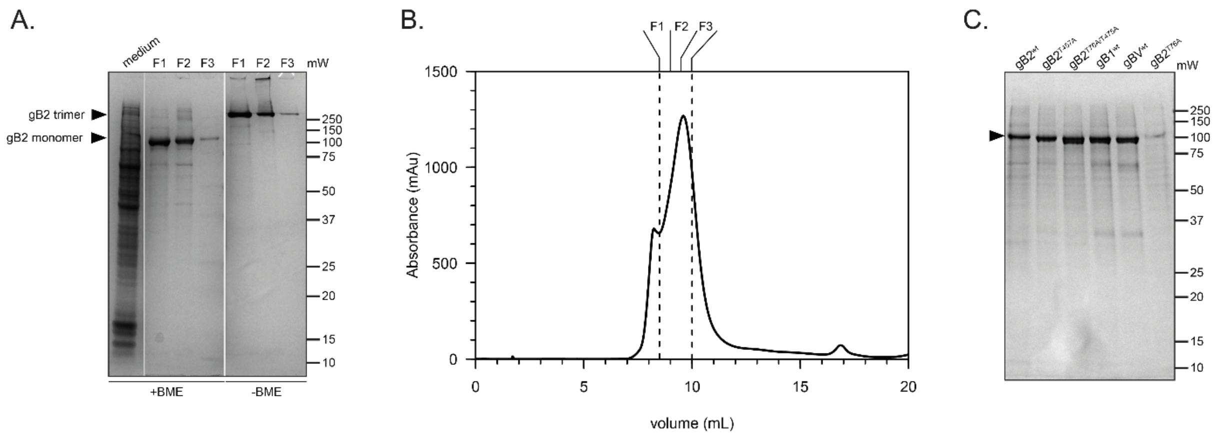


**Supplementary Figure S6.** Purification of gB2 and gB constructs. A. 12% SDS-PAGE gel of a large scale purification of gB2. Medium supernatant sample was taken before affinity column loading. Size Exclusion Chromatography (SEC) fractions (F1-3) were loaded on the gel to assess purity. BME = β-mercaptoethanol added to the loading dye. Expected band locations based on trimeric and monomeric gB2 calculated molecular weight are indicated by black triangles. B. UV280 absorption SEC profile for gB2. A superdex200 Increase column (Cytiva) was used for separation. Fractions F1-3 are indicated. C. Small scale purification of gB2 mutants, gB1 and gBV for SPR. Elutions of affinity purifications were loaded on a reducing (+BME) SDS-PAGE gel. Expected size for all constructs is indicated by a black triangle.

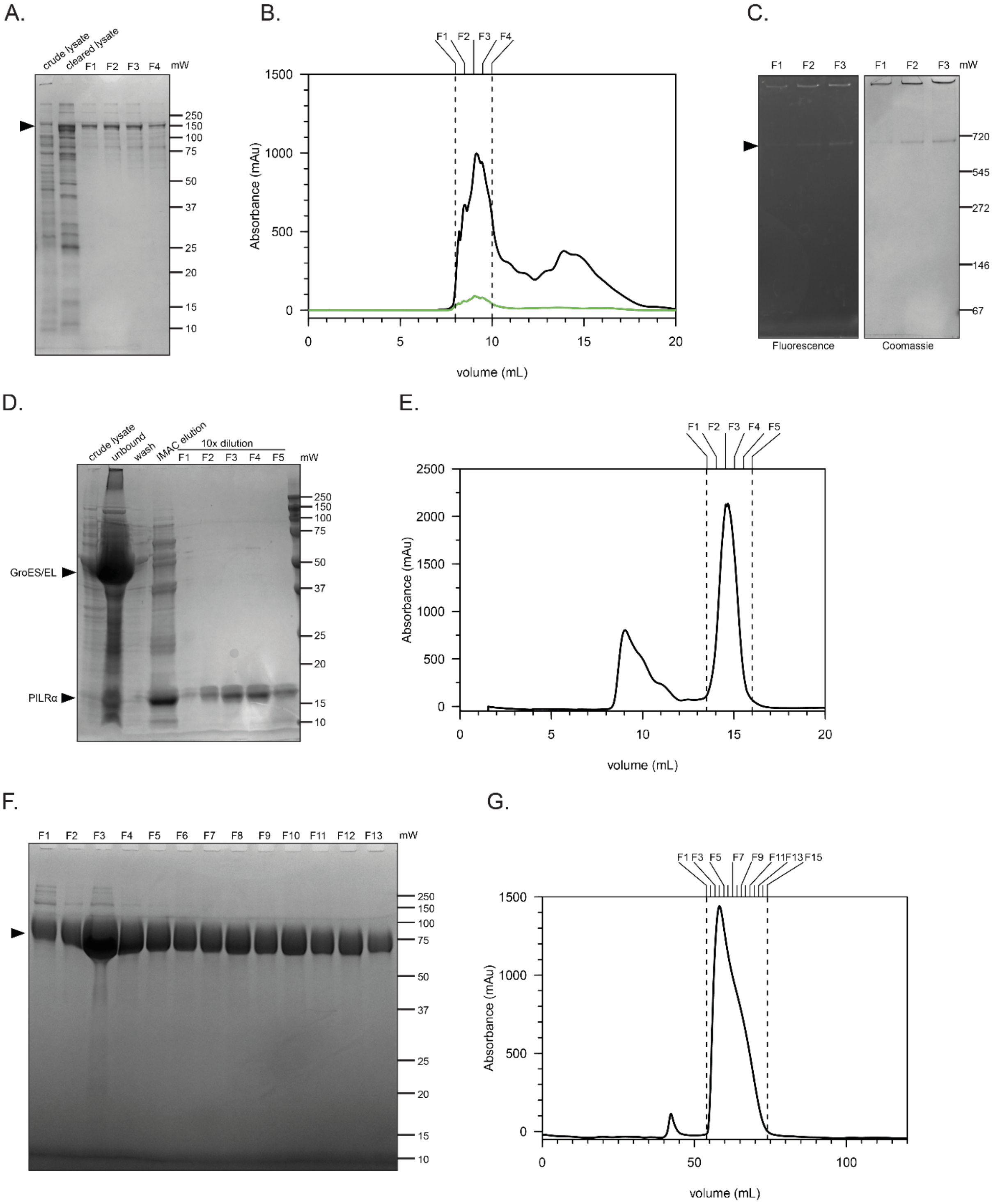


**Supplementary Figure S7.** Purification of ectodomains of gB co-receptors. A-C. Purification of NM-IIA coiled coil domain with N-terminal GFP fusion. A. 12% reducing SDS-PAGE gel of NM-IIA samples. Sample of E.coli cells + medium before lysis is indicated by "crude lysate", supernatant sample taken after lysis and ultracentrifugation is indicated by "cleared lysate". Four Size Exclusion Chromatography (SEC) fractions (F1-4) are loaded on the gel to assess purity. Expected NM-IIA construct molecular weight is indicated with a black triangle. B. UV280 and UV501 (GFP) absorption SEC profiles of NM-IIA using a Superose 6 Increase column (Cytiva). F1-4 are indicated. C. GFP fluorescence signal of purified NM-IIA fractions on a 4-12% native SDS-PAGE gradient gel. GFP fluorescence was imaged in cartridge before gel staining. D and E. Purification of PILRα. D. 12% reducing SDS-PAGE gel of PILRα purification. Sample of E.coli cells + medium before lysis is indicated by "crude lysate", flowthrough sample taken after affinity column loading is indicated by "unbound", flowthrough sample of buffer wash is indicated by "wash" and IMAC elution sample by "IMAC elution". SEC fractions (F1-5) were diluted 1:10 before loading. E. UV280 absorption SEC profile of PILRα. Fractions F1-5 are indicated. A superdex75 Increase column (Cytiva) was used. F-G. Purification of MAG. F. 12% reducing SDS-PAGE gel of SEC fractions (F1-13) of MAG. Expected molecular weight of MAG is indicated by a black triangle. G. UV280 absorption SEC profile of MAG. Fractions 1-13 are indicated. A HiLoad 16/60 s200 increase column (Cytiva) was used.
